# Evolutionarily Conserved *Plasmodium falciparum* HSP70-3 Enhances the Immunogenicity of the Established Malaria Vaccine Antigen Circumsporozoite Protein through Cross-Presentation

**DOI:** 10.64898/2026.10.15.759406

**Authors:** Prachi Garg, Chandan Kumar Verma, Kanika Gupta, Ayushee Sharma, Shivam Jawla, Neha Jha, Savita Saini, Kota Arun Kumar, Anand Ranganathan, Shailja Singh

## Abstract

**Background:** Clinically approved *P. falciparum* circumsporozoite protein (*Pf*CSP)-based pre-erythrocytic stage malaria vaccines suffer from sub-optimal efficacy and rapidly waning memory. By decoding these limitations, we fast-tracked vaccine optimization by correcting CSP’s fragile immunogenicity using a parasite-derived chaperone. Unlike other orthologues of *P. falciparum* heat shock protein 70 (*Pf*HSP70), substrate-binding domain (SBD) of *Pf*HSP70-3 is an evolutionarily conserved internal mitochondrial matrix chaperone. We demonstrate that *Pf*HSP70-3-SBD hydrophobic region complexes with *Pf*CSP and targets host CD91 on antigen-presenting cells to trigger cross-presentation, resulting in enhanced immune responses to *Pf*CSP.

**Methods:** *In silico* analyses predicted interactions among *Pf*HSP70-3-SBD, *Pf*CSP, and CD91 receptor. Recombinant *Pf*HSP70-3-SBD and *Pf*CSP proteins were purified and assessed for *in vitro* interaction using ELISA and far-western dot blot assays. BALB/c mice were immunized with alum-formulated *Pf*CSP, with or without *Pf*HSP70-3-SBD. *Pf*CSP-specific IgG and IFN-γ expression were evaluated by ELISA and RT-qPCR, respectively. Functional activity was assessed using transgenic *Plasmodium berghei* expressing *Pf*CSP (*Pb/Pf* CSP), by sporozoite neutralization assay.

**Findings:** *Pf*HSP70-3-SBD demonstrated a specific interaction with *Pf*CSP. The complex primarily engages CD91 receptor cluster IV, providing a potential mechanistic basis for antigen cross-presentation. Incorporation of *Pf*HSP70-3-SBD enhanced *Pf*CSP-specific total IgG responses and increased IFN-γ expression in BALB/c mice, indicating improved immunogenicity upon co-formulation. Notably, immune sera neutralized *Pb/Pf* CSP sporozoites, delaying blood-stage patency until day 12 in 40% of C57BL/6 mice, with no detectable parasitemia in surviving mice for up to 2 months, indicating sustained efficacy against the sporozoite stage.

**Interpretation:** Molecular chaperones, including HSPs, are currently under intense investigation for their immune-potentiating activity, leading to next-generation vaccine development. By utilizing chaperonic domains derived directly from the pathogenic agent itself to serve an essential adjuvant function, this approach reduces reliance on heterologous adjuvants while enhancing antigen-specific immunity.

**Funding:** This work was supported by funding from Indian Council of Medical Research (IIRPSG-2024-01-03940).

**RESEARCH IN CONTEXT:** 

**Evidence before this study:** Evidence from *Pf*CSP-based licensed vaccines, RTS,S/AS01 and R21/Matrix-M, underscores the need to improve the magnitude and durability of vaccine-induced protection. Although alum is an established adjuvant used in human vaccines, its limited capacity to elicit robust Th1-type cellular responses motivates the search for complementary strategies to fight against malaria. Findings on *P. falciparum* heat shock proteins (HSPs) remain mixed. *Pf*HSP70 has been reported to lack immunomodulatory activity, whereas an HSP70-derived C-terminal fragment enhanced DNA vaccine potency when linked to a malaria antigen. Whether mitochondrial *Pf*HSP70-3 substrate-binding domain can enhance *Pf*CSP immunogenicity and function as adjuvant remains insufficiently characterized. Although CD91-mediated uptake and cross-presentation of HSP-bound antigens have been previously reported, the potential involvement of this pathway in *Pf*CSP-*Pf*HSP70-3-SBD complex-mediated antigen presentation remains unexplored.

**Added value of this study:** This study explores *Pf*HSP70-3-SBD as a novel parasite-derived adjuvant, offering an alternative approach to fast-track the improvement of *Pf*CSP-based subunit vaccine immunogenicity. By linking antigen-chaperone interactions with the potential involvement of host CD91 receptor, it provides a plausible mechanistic framework for investigating enhancement in *Pf*CSP antigen cross-presentation. The integration of immunological evaluation with a functional neutralizing activity against transgenic *P. berghei* sporozoites expressing *P. falciparum* CSP, extends the findings beyond immunogenicity.

**Implications of all the available evidence:** *Pf*HSP70-3-SBD represents a promising candidate for further development as a parasite-derived chaperonic domain with modular adjuvant activity for next-generation malaria subunit vaccines. Incorporating *Pf*HSP70-3-SBD into *Pf*CSP-based formulations may offer a complementary approach to existing adjuvant strategies while providing a basis for investigating antigen-specific cross-presentation. This strategy may ultimately facilitate more streamlined vaccine development by integrating antigen delivery with immune potentiation. This approach further represents an initial step towards developing parasite-derived adjuvants, to reduce reliance on heterologous adjuvant sources.

## INTRODUCTION

The first line of treatment for malaria infections is currently artemisinin (ART) and ART-based combination treatments (ACTs). However, with the increasing ART resistance in various areas, vaccination against malaria has become a major prevention strategy to be applied prior or concomitantly with therapeutic development. Recently, there has been a significant focus on ongoing preclinical and clinical trials aimed at developing an efficacious malaria vaccine. Among various *P. falciparum* antigens, circumsporozoite protein (*Pf*CSP) remains the leading pre-erythrocytic stage candidate for vaccine development. Since *Pf*CSP is a major sporozoite-stage surface protein that mediates parasite invasion into hepatocytes, it represents an attractive vaccine target for blocking liver-stage infection and preventing the establishment of malaria.^1–3^ Till now, two malaria vaccines, RTS,S/AS01 and R21/Matrix-M, which target the *Pf*CSP antigen during the pre-erythrocytic stage of the parasite life cycle, have been recommended by the World Health Organization (WHO).^4,5^ Despite the deployment of licensed malaria vaccines, protection remains incomplete and may decline over time. This highlights an urgent need for next-generation vaccine formulations that can induce stronger, more durable and efficacious immune responses.

Most of the traditional vaccine strategies used routinely against *P. falciparum* are protein subunit vaccines, which are inherently safe but poorly immunogenic on their own.^6^ Therefore, subunit vaccine antigens require potent adjuvants to generate robust immunogenicity with translational potential for human application. One extensively used adjuvant in human vaccines is alum that predominantly induces a Th2-biased immune response and effectively enhances humoral immunity, but possesses limited ability to stimulate a Th1-type cellular immune response.^7–9^ However, CD8^+^ T cells contribute substantially to achieve protective immunity against the liver stage of the parasite life cycle.^10^ Consequently, substantial efforts have been focused on enhancing vaccine immunogenicity through the incorporation of novel adjuvants together with targeted antigen delivery with the aim to simultaneously elicit long-lasting cellular responses in conjunction with robust humoral immunity.

In this regard, heat shock proteins (HSPs) have become attractive immunostimulating agents for vaccine development against various infectious diseases and cancers. HSPs are molecular chaperones that are induced upon physical stresses and play a vital role in cellular proteostasis, thereby providing a robust protein quality control mechanism.^11^ The two distinct functional domains of HSP70s are the highly conserved N-terminal ATPase domain and the less conserved C-terminal substrate-binding domain (SBD), which are joined by a short linker.^12^ These molecular chaperones are known to amplify both innate and antigen-specific effector functions.^13^ Their immunostimulatory activity is attributed to their capacity to activate dendritic cells, leading to secretion of proinflammatory cytokines and chemokines along with enhanced antigen presentation.^14^ HSPs possess a wide range of peptide and protein binding capabilities. This characteristic is shared with major histocompatibility complex (MHC) molecules and is fundamental to the presentation of antigens to the adaptive immune system.^15–18^ This property enables exogenous antigens chaperoned by HSPs to be presented by MHC class I molecules and subsequently recognized by CD8^+^ T lymphocytes.^19,20^ Together with their capacity to induce antigen-specific CD4⁺ T-cell responses through the MHC class II pathway, these properties support the potential of HSPs as effective molecular adjuvants for protein subunit vaccines.^13,21^

A plethora of pathogens have been reported with the presence of heat shock proteins and are suggested to possess adjuvant properties for the development of novel vaccines requiring both humoral and cell-mediated immunity against infectious diseases. Molecular chaperones constitute approximately 2% of the *Plasmodium falciparum* genome for structural integrity of the proteome that contributes to the survival and virulence of malaria parasite.^22^ *P. falciparum* HSP70 (*Pf*HSP70) represents one of the major molecular chaperone families, strongly adapted to maintain the proteome’s structural and functional integrity. *P. falciparum* has six orthologues of HSP70s, including *Pf*HSP70-1, *Pf*HSP70-2, *Pf*HSP70-3, *Pf*HSP70-x, *Pf*HSP70-y, and *Pf*HSP70-z, that are finely tuned to the parasite lifecycle.^23^ All six orthologues of *Pf*HSP70s possess the classical domains but differ in their subcellular localization within the parasite.^24,25^ Although several members show anti-malarial activity, accumulating evidence indicates the adjuvant effect of a C-terminal region of *Pf*HSP70, which enhanced the potency of a DNA vaccine by genetic linkage with *P. falciparum* antigen.^26^

Among the HSP70 family members of *P. falciparum*, *Pf*HSP70-3 is uniquely predicted to localize to the parasite mitochondrion, making it a distinct and relatively underexplored member of the parasite HSP70 repertoire. Importantly, *Pf*HSP70-3 retains conserved structural elements of the HSP70 substrate-binding cavity, including the conserved valine residue within the hydrophobic substrate-binding pocket, which has been implicated in substrate access.^24,27,28^ This conserved feature suggests that *Pf*HSP70-3 possesses a structurally competent substrate-binding cavity, despite its distinct subcellular localization and sequence divergence. Structurally, the substrate-binding domain of HSP70s comprises the SBDβ and SBDα subdomains, of which SBDβ forms the principal substrate-binding interface, while SBDα functions as a lid that stabilizes the bound peptide.^29–31^. Thus, the peptide-binding capacity of SBDβ provides a rational structural basis for exploring this domain beyond its canonical chaperone function, particularly as an antigen-interacting module capable of facilitating antigen delivery and subsequent cross-presentation. Based on this rationale, we selected the β-subdomain of the *Pf*HSP70-3-SBD for evaluation of its potential immunopotentiating properties.

In this study, we evaluated whether co-formulation of the recombinant *Pf*HSP70-3-SBD with a leading pre-erythrocytic stage antigen, *Pf*CSP, enhances antigen-specific immunity in a murine model. In this regard, we first demonstrate that *Pf*HSP70-3-SBD interacts with *Pf*CSP using *in silico* protein-protein docking studies. To confirm their *in vitro* interaction, we next expressed and purified hexahistidine-tagged recombinant *Pf*CSP and *Pf*HSP70-3-SBD proteins in *E. coli* with high purity and demonstrated a strong interaction between the two proteins, supporting the potential role of *Pf*HSP70-3-SBD in enhancing antigen immunogenicity. For preclinical evaluation, BALB/c mice were immunized with *Pf*CSP/Alum with or without *Pf*HSP70-3-SBD. Incorporation of *Pf*HSP70-3-SBD into the *Pf*CSP/Alum vaccine formulation consistently augmented antigen-specific humoral and cellular immune responses as studied through antigen-specific total IgG responses and qRT-PCR assays. To evaluate the functional activity of immune sera, a transgenic *P. bergehi* parasite line expressing *P. falciparum* CSP (*Pb/Pf* CSP) was employed. Interestingly, functional evaluation of vaccine-induced immune sera demonstrated the sporozoite-neutralizing potential of the induced antibodies. Collectively, these findings position *Pf*HSP70-3-SBD as a promising chaperonic domain with adjuvant-like activity for malaria subunit vaccine development. This study further establishes *Pf*HSP70-3-SBD as an adjuvant-like domain capable of strengthening antigen-specific immune responses and provides a conceptual framework for repurposing parasite-derived chaperonic domains as rational immune adjuvant candidates for next-generation malaria subunit vaccines.

## METHODS

### Bacteria and media

The *Escherichia coli* BL21(DE3) expression strain was employed for recombinant protein expression and purification. Luria-Bertani (LB) agar (HiMedia, Cat. No. M1151-500G) plates were incubated at 37 °C, and LB broth (HiMedia, Cat. No. M1245-500G) cultures were maintained at 37 °C with continuous shaking. For the selection of strains harbouring the recombinant pET-28(b)+ plasmid, the medium was supplemented with kanamycin (HiMedia, Cat. No. CMS210-5G) to a final concentration of 50 μg/mL. Streptomycin (AMRESCO, Cat. No. 0382-100G) was used at 50 μg/mL for the selection of strains containing the recombinant pMTSAra plasmid.

### Parasite propagation, mosquito infection and sporozoite preparation

To produce *P. berghei* sporozoites expressing *P. falciparum* CSP, (*Pb/Pf* CSP),^32^ female BALB/c mice were used for asexual propagation of the transgenic parasites. Malaria transmission to mosquito was achieved by allowing female *Anopheles stephensi* mosquitoes to obtain two consecutive blood meals from anaesthetized BALB/c that carried 1-2% gametocytemia. The blood fed mosquitoes were maintained at 20 °C and 80% of relative humidity. The midgut load of the parasite was assessed on day 14 post-blood meal by examining the presence of midgut oocysts under a light microscope. Further, salivary glands were dissected on day 20 post-blood meal in incomplete RPMI-1640 (Gibco, Cat. No. 22400-071) medium. The dissected glands were collected in 1.5 ml Eppendorf tube and mechanically disrupted using plastic pestle. The sample was centrifuged at 0.06 x g (800 rpm) for 3 min at 4 °C. The salivary gland sporozoites were enumerated using a hemocytometer.

### *In silico* protein-protein docking

The domain boundaries for *Pf*CSP (PlasmoDB ID: *PF*3D7_0304600) were assigned based on US Patent 11,896,655 B2 and corroborated with existing literature.^33–35^ Domain boundaries for *Pf*HSP70-3 (PlasmoDB ID: *PF*3D7_1134000) were assigned based on the InterPro domain annotation for UniProt accession Q8II24, together with the reported mitochondrial transit peptide boundary.^36,37^ In this study, we selected the *Pf*CSP region (179 residues, amino acids 199-377) encompassing the central NANP repeat and C-terminal regions, and the β-subdomain of the substrate-binding domain of *Pf*HSP70-3 (113 residues, amino acids 427-539) was selected. The three-dimensional structures for *Pf*CSP and *Pf*HSP70-3-SBD, along with *Homo sapiens* CD91 receptor (UniProt accession Q07954) for Cluster II (*Hs*CD91-CLII, 380 residues, amino acids 786-1165) and Cluster IV (*Hs*CD91-CLIV, 448 residues, amino acids 3332-3779),^38^ were generated using AlphaFold3 server,^39^ and the quality of the structure for its stereochemical properties was assessed using ERRAT,^40^ and PROCHECK.^41^ To further check the accuracy of the AlphaFold3 predicted structure of *Pf*CSP, the properly folded region comprising the NANP repeats and C-terminal domain was aligned and compared to the crystal structure provided for the *Pf*CSP α-thrombospondin type-1 repeat (αTSR) domain (PDB ID: 3VDJ).^42^ The AlphaFold3 predicted model for *Pf*HSP70-3-SBD was also pairwise aligned and compared with the crystal structure of the substrate-binding domain of the orthologous *Pf*HSP70-x (PDB ID: 6ZHI),^43^ whose Cα root mean square deviation (RMSD) was observed by super positioning both the structures using PyMOL.^44^

To systematically evaluate the interaction of *Pf*CSP-*Pf*HSP70-3-SBD, we performed molecular docking using three complementary methods. First, a rigid-body-based blind docking using HDOCK,^45,46^ in which we used the *Pf*CSP region along with *Pf*HSP70-3-SBD without prior restraints. Second was the information-driven flexible docking, which was performed in HADDOCK 2.4,^47^ where we used Ambiguous Interaction Restraints (AIRs), in which active and passive residues were derived from the AlphaFold3-predicted interface for both proteins. The resulting docking poses were then clustered based on default RMSD-based clustering in HADDOCK, and the pose having the best HADDOCK score and Z-score was selected. Third was the complementary docking, in which we used both docking protocols with the crystal structure of the *Pf*CSP αTSR domain (PDB ID: 3VDJ) and *Pf*HSP70-x (PDB ID: 6ZHI) instead of the AlphaFold3-predicted models, to independently validate the predicted interaction interface. Interface residues were defined using the RING 2.0 web server,^48^ and for estimation of the binding affinity of the top-ranked model, we used the PRODIGY web server.^49^ To further evaluate the interaction of *Pf*CSP-HSP70-SBD complex with host *Hs*CD91-CLII and *Hs*CD91-CLIV required for cross-presentation by APCs, molecular docking was performed using HDOCK, and ClusPro,^50^ and interface residues were defined using the RING 2.0 web server.

### Expression and protein purification of substrate-binding domain of *Pf*HSP70-3 (*Pf*HSP70-3-SBD)

The gene encoding the substrate-binding domain β-sandwich subdomain of *Pf*HSP70-3 was synthesized into the pMTSAra vector between the SnaBI restriction enzyme sites after being codon-optimized for effective expression in *E. coli*. The amino acid sequence details are provided in **Table S1.** The recombinant pMTSAra-*Pf*HSP70-3-SBD vector was transformed into competent *E. coli* BL21 (DE3) cells. This was followed by the selection of transformants using streptomycin (50 μg/mL) for recombinant protein expression and purification. BL21 cells carrying the pMTSAra-*Pf*HSP70-3-SBD plasmid were cultured overnight in LB broth supplemented with streptomycin at 37 °C. Next day, secondary culture was inoculated using 1% inoculum from primary cultures. After being grown until the mid-log phase, cultures were induced with 0.4% arabinose (SRL Pvt. Ltd., Cat. No. 52392) for five hours at 37 °C. Cell pellet was resuspended in lysis buffer containing 50 mM Tris pH 8.0, 100 µg/ml lysozyme, 10 mM EDTA, and 2 mM PMSF and incubated at 4 °C for 1 hour. After that, cells were lysed by sonication, and a clear cell lysate and pellet were obtained by centrifugation. Since the protein was recovered predominantly in inclusion bodies, pellet was washed using pellet washing buffer (50 mM Tris pH 8.0, 100 mM NaCl, 0.5% Triton X100, and 0.1% sodium azide). Solubilization was done using solubilization buffer (8 M urea, 50 mM Tris, 100 mM NaCl, pH 8.0) and incubated for complete solubilization. The clear suspension obtained after centrifugation was incubated with Ni-NTA (QIAGEN, Cat. No. 30230) beads for efficient binding at room temperature (RT) for 2 hours. The elution of bound protein was done with an affinity column by gradually increasing the imidazole (Sigma-Aldrich, Cat. No. 56750-100G) concentration in the solubilization buffer. Eluted protein fractions were run on a 15% sodium dodecyl sulfate-polyacrylamide gel electrophoresis (SDS-PAGE). Pure fractions with high protein concentration were pooled and refolded overnight at 4 °C in a buffer containing 100 mM Tris, 20 % glycerol, 250 mM L-arginine, 1 mM EDTA, pH 8.0, along with 1 mM reduced glutathione and 0.5 mM glutathione disulfide. Dialysis was performed at 4 °C with dialysis buffer (10 mM HEPES, 150 mM NaCl, pH 8.2). SDS-PAGE was used to confirm the size and integrity of recombinant protein. Further, coomassie brilliant blue (CBB) staining and western blotting using anti-His antibody was performed.

### Expression and purification of *P. falciparum* CSP (*Pf*CSP) region comprising NANP repeats and C-terminal regions

For cloning of the *Pf*CSP region corresponding to the central NANP repeat and C-terminal regions, the codon-optimized gene sequence was cloned in pET-28(b)+ vector using EcoRI (Thermo Scientific, Cat. No. FD0274) and HindIII (Thermo Scientific, Cat. No. FD0504) restriction enzymes. The amino acid sequence details are provided in **Table S1**. Competent *E. Coli* BL21 (DE3) cells were transformed with the recombinant plasmid. Positive clones were selected using kanamycin (50 µg/mL). Primary cultures were grown overnight in LB broth with kanamycin at 37 °C. Secondary cultures were grown to mid log phase and protein induction was carried out using 1 mM isopropyl β-D-1-thiogalactopyranoside (SRL Pvt. Ltd., Cat. No. 67208) for 5 hours at 37 °C. The culture pellet was resuspended in lysis buffer (20 mM Tris pH 8.0, 150 mM NaCl, 2 mM PMSF, 100 µg/mL lysozyme) and incubated at 4 °C for 1 hour. Cell lysis was performed by sonication and centrifugation was done to obtain clear supernatant and pellet. Recombinant *Pf*CSP protein was obtained in the supernatant fraction. The supernatant was incubated with Ni-NTA beads for 2 hours at 4 °C. Using increasing imidazole concentrations, protein was subsequently eluted using affinity column in 20 mM Tris pH 8.0, 150 mM NaCl. 12% SDS-PAGE was used to run the eluted fractions, and the fractions with high purity and protein concentration were pooled and subjected to dialysis against 10 mM Tris buffer pH 8.0. Subsequently, PBS (pH 7.4) was used to carry out buffer exchange. The integrity and size of the protein were confirmed on SDS-PAGE followed by CBB-staining and western blotting using anti-His antibody.

### Western blot analysis

Recombinant protein or transgenic *Pb/Pf* CSP sporozoite lysates were suspended in Laemmli buffer and loaded on SDS-PAGE. Proteins were transferred to polyvinylidene difluoride (PVDF) transfer membranes (mdi Membrane Technologies, Cat. No. SVFX8301XXXX101). 5% skimmed milk (SRL Pvt. Ltd., Cat. No. 28582) in 1× PBS buffer was added to block the membranes overnight at 4 °C. Next day, membranes were washed thrice with PBST buffer (1× PBS containing 0.05% Tween 20). At room temperature, primary antibodies were added and incubated for two hours. PBST was used to wash the membranes three times and subsequently incubated with secondary antibodies conjugated with HRP for 1 hour at room temperature. After stringent washing, enhanced chemiluminescence (ECL) kit (Bio-Rad, Cat. No. 1705060) was used for visualization of blots on the Bio-Rad ChemiDoc Imaging System.

Mouse anti-His antibody (Elabscience, Cat. No. E-AB-20009) was used as primary antibody for histidine-tagged recombinant proteins at a dilution of 1:10,000. Subsequently, secondary anti-mouse antibody conjugated with HRP (Sigma-Aldrich, Cat. No. A9044) was added at 1:5,000 dilution. In a separate experiment, vaccine-induced sera were assessed for the recognition of *Pf*CSP antigen from transgenic *Pb/Pf* CSP salivary gland sporozoite lysates. Anti-*Pf*CSP/*Pf*HSP70-3-SBD/Alum immune sera were assessed for recognition of *Pf*CSP and thereby utilized as primary antibody. HRP-conjugated anti-mouse antibody (Cell Signaling Technology, Cat. No. 7076S) was added subsequently as secondary antibody at a dilution of 1:5,000.

### ELISA-based protein-protein interaction assay

To validate the interaction of *Pf*HSP70-3-SBD and *Pf*CSP, ELISA was performed. 200 ng of recombinant *Pf*HSP70-3-SBD protein (bait) was coated per well using coating buffer (BioLegend, Cat. No. 421701) in a flat-bottom 96 well MaxiSorp ELISA plate (Thermo Fisher Scientific, Cat. No. 442404). Coating of *Pf*HSP70-3-SBD protein was done overnight at 4 °C. The next day, blocking was done at 37 °C for 1 hour using 5 % bovine serum albumin (BSA, SRL Pvt. Ltd., Cat. No. 85171) prepared in PBST. Plates were washed with PBST and incubated for 2 hours at 37 °C with serially diluted concentrations of *Pf*CSP (prey) ranging from 31.25 ng to 2000 ng, as an interacting partner. Following stringent washes with PBST buffer, plates were incubated with rabbit monoclonal antibody 2A10 (Absolute Antibody, Cat. No. Ab03231-23.0) at a dilution of 1:3,000 (prepared in 1% BSA in PBST buffer) for 2 hours at 37 °C. After washing, plates were incubated with HRP-conjugated anti-rabbit secondary antibody (Sigma-Aldrich, Cat. No. A9169) at a dilution of 1:8,000 for 1 hour at 37 °C. Plates were washed, and 100 μl of 3,3′,5,5′-tetramethylbenzidine (TMB) substrate (Thermo Scientific, Cat. No. 34021) was used for developing the plates. The reaction was terminated using 2N sulphuric acid. Using Varioskan LUX Multimode Microplate Reader (Thermo Scientific), absorbance was measured at 450 nm. Interaction between *Pf*CSP and BSA served as negative control during the experiment.

### Far-Western dot blot assay

For protein-protein interaction study, a far-western dot blot assay was performed. 2.5 μg each of *Pf*HSP70-3-SBD and BSA were immobilized onto a nitrocellulose membrane (Bio-Rad, Cat. No. 1620112). Next, the membrane was allowed to dry for 30 minutes and blocked overnight at 4 °C using 5 % BSA prepared in 1×PBS. The next day, the membrane was washed three times with PBST wash buffer and incubated with 5 μg/mL of *Pf*CSP as secondary protein in 1% BSA in PBST for 2 hours at RT. Unbound secondary protein was removed by thoroughly washing the membrane three times with PBST. Further, membranes were incubated with rabbit mAb 2A10 (specific for *Pf*CSP repeats) at 1: 1,000 dilution for 2 hours at RT. After 2 hours, the membrane was washed with PBST and incubated with HRP-conjugated anti-rabbit secondary antibody (1:5,000; Sigma-Aldrich, Cat. No. A9169) for 1 hour at RT. Similarly, in a separate reciprocal experiment, 2.5 μg each of *Pf*CSP and BSA were immobilized on the membrane and 5 μg/ml of *Pf*HSP70-3-SBD was added as secondary protein. Membrane was incubated with in-house-generated polyclonal mouse anti-*Pf*HSP70-3-SBD sera at a dilution of 1:1,000. Further, HRP-conjugated anti-mouse secondary antibody (1:5,000; Sigma-Aldrich, Cat. No. A9044) was added. ECL kit (Bio-Rad, Cat. No. 1705060) was used for the visualization of blots on the Bio-Rad ChemiDoc Imaging System. Densitometry analysis was performed using ImageJ software and arbitrary densitometry units (ADUs) were calculated.

### Immunization of mice

Male BALB/c mice aged four to six weeks were obtained from Central Laboratory Animal Resources (CLAR), Jawaharlal Nehru University (JNU), New Delhi. Mice were allocated into three experimental groups (n=5 per group). For immunization, each BALB/c mouse was injected intramuscularly with 40 µg of recombinant *Pf*CSP antigen formulated in alum (G-Biosciences, Cat. No. 786-1216) with or without co-formulation with 40 µg of recombinant *Pf*HSP70-3-SBD protein. Formulations were prepared in PBS, with a total volume of 100 µl used for immunization of each mouse. Immunizations were performed on day 0 (prime-dose), 14 (first booster-dose), and 28 (second booster-dose). A group of mice was injected intramuscularly with only PBS formulated in alum adjuvant as a control. Blood was collected by retro-orbital plexus puncture on days 13, 27, and 42. Sera were stored at -20 °C for downstream immunological and neutralization studies. On day 42, mice were euthanized, and their spleens were aseptically removed and collected for further immunological assays. Animal experimentation procedures were conducted in compliance with the Institutional Animal Ethics Committee (IAEC) of Jawaharlal Nehru University and University of Hyderabad.

### IgG titer determination by ELISA

The levels of total IgG antibodies against *Pf*CSP were quantified using an indirect ELISA. Briefly, flat-bottom 96-well MaxiSorp ELISA plates (Thermo Fisher Scientific, Cat. No. 442404) were coated overnight at 4 °C with 200 ng of recombinant *Pf*CSP per well in coating buffer (BioLegend, Cat. No. 421701). Next day, plates were washed thrice with PBST and blocked for 1 hour at 37 °C with 5% BSA prepared in PBS. Following three washes with PBST, serum samples were added to the antigen-coated wells for incubation at 37 °C for 2 hours. Mouse serum samples were serially diluted in 1% BSA-containing PBST, covering dilutions ranging from 1:5,000 to 1:40,000. After three washes with PBST, wells were incubated for 1 hour at 37 °C with an HRP-labeled anti-mouse IgG secondary antibody (1:8,000; Sigma-Aldrich, Cat. No. A9044). Following the final washing step, plates were developed using TMB substrate. To stop the reaction, 2 N sulfuric acid (H₂SO₄) was added, and optical density was recorded at 450 nm using Varioskan LUX Multimode Microplate Reader (Thermo Scientific).

### Quantitative real-time PCR

Total RNA was isolated from the whole spleen of mice fourteen days after the second booster immunization. RNA isolation was carried out using TRIzol reagent (Invitrogen, Cat. No. 15596026) in RNase-free water. cDNA was synthesized from 1 µg of total RNA using the iScript cDNA Synthesis Kit (Bio-Rad, Cat. No. 1708891), according to the manufacturer’s protocol. Real-time PCR (RT-PCR) was performed using a Real-Time PCR System (Applied Biosystems) with SYBR Green Universal Master Mix (Applied Biosystems, Cat. No. 43-091-55). IFN-γ levels were quantified using specific forward and reverse primers. Mouse GAPDH amplification was used as an internal control for normalization. Primer sequences are provided in **Table S2.** The temperature cycling conditions were as follows: 50 °C for 2 minutes, then 40 cycles of 95 °C for 15 seconds, 60 °C for 30 seconds, and 72 °C for 1 minute. The 2^−ΔΔCt^ method was used to measure gene expression levels, which were normalized to the GAPDH housekeeping gene. GraphPad Prism version 8.0 was used to plot and analyze the data.

### Transgenic sporozoite neutralization assay (TSNA)

Vaccine-induced neutralizing antibodies were tested for their functional inhibitory activity using a transgenic *Plasmodium berghei* parasite line expressing *Plasmodium falciparum* circumsporozoite protein (CSP) repeats (*Pb/Pf* CSP).^32^ Approximately, 5 × 10³ partially purified *Pb/Pf* CSP sporozoites were incubated with immune sera at 1:10 dilution, for 35 minutes on ice. The neutralized sporozoites were immediately injected via intravenous route naive C57BL/6 mice (n=5/group), aged 9-10 weeks old (purchased from Hylasco Bio-Technology (India) Private Limited). Control groups consisted of sporozoites incubated with sera from unimmunized mice (1:10) or with monoclonal antibody 2A10 (1:50 dilution of 1 mg/mL antibody concentration, used as positive control), which specifically targets *P. falciparum* CSP repeats. From day 3.5 post-infection with sporozoites, Giemsa-stained thin blood smears were prepared, and the pre-patent period was determined, till day 12 post-infection. Furthermore, surviving mice were monitored for 2 months after sporozoite infection to assess long-term outcomes. All procedures were conducted in accordance with the Institutional Animal Ethics Committee (IAEC) of the University of Hyderabad.

### Statistical analyses

GraphPad Prism version 8.0 was used for plotting the data. Means ± standard deviation (SD) were used to express the data. The differences between the experimental groups were assessed and compared using an unpaired Student’s t-test and p-values are displayed. In TSNA, Kaplan-Meier curves were generated and for statistical analysis, Log-rank (Mantel-Cox) test was used.

## RESULTS

### *In silico* docking predicts an interaction between the *Pf*HSP70-3 substrate-binding domain β-subdomain and the *P. falciparum* CSP region

The domain architecture of *Pf*CSP (PlasmoDB ID: *PF*3D7_0304600), with the central repeat region and TSR domain used for docking, is represented by lime green box (**Fig. 1A (i)),** and structural model of *Pf*CSP region; was generated using AlphaFold3 (**Fig. 1A (ii))**. Model quality was assessed for its stereochemical properties using the ERRAT and PROCHECK web servers, and 3Drefine was used for refinement and energy minimization; ERRAT showed an overall quality factor of 100%, with 99.3% of residues falling in the Ramachandran favoured region **(Fig. S1)**. To validate the accuracy of this model, the folded TSR domain was superposed onto the crystallographically solved *Pf*CSP αTSR domain (PDB ID: 3VDJ), yielding a Cα RMSD of 0.315 Å (**Fig. 1A (iii))**, confirming that AlphaFold3 accurately reproduced the experimentally observed fold of this region.

**Fig. 1:**
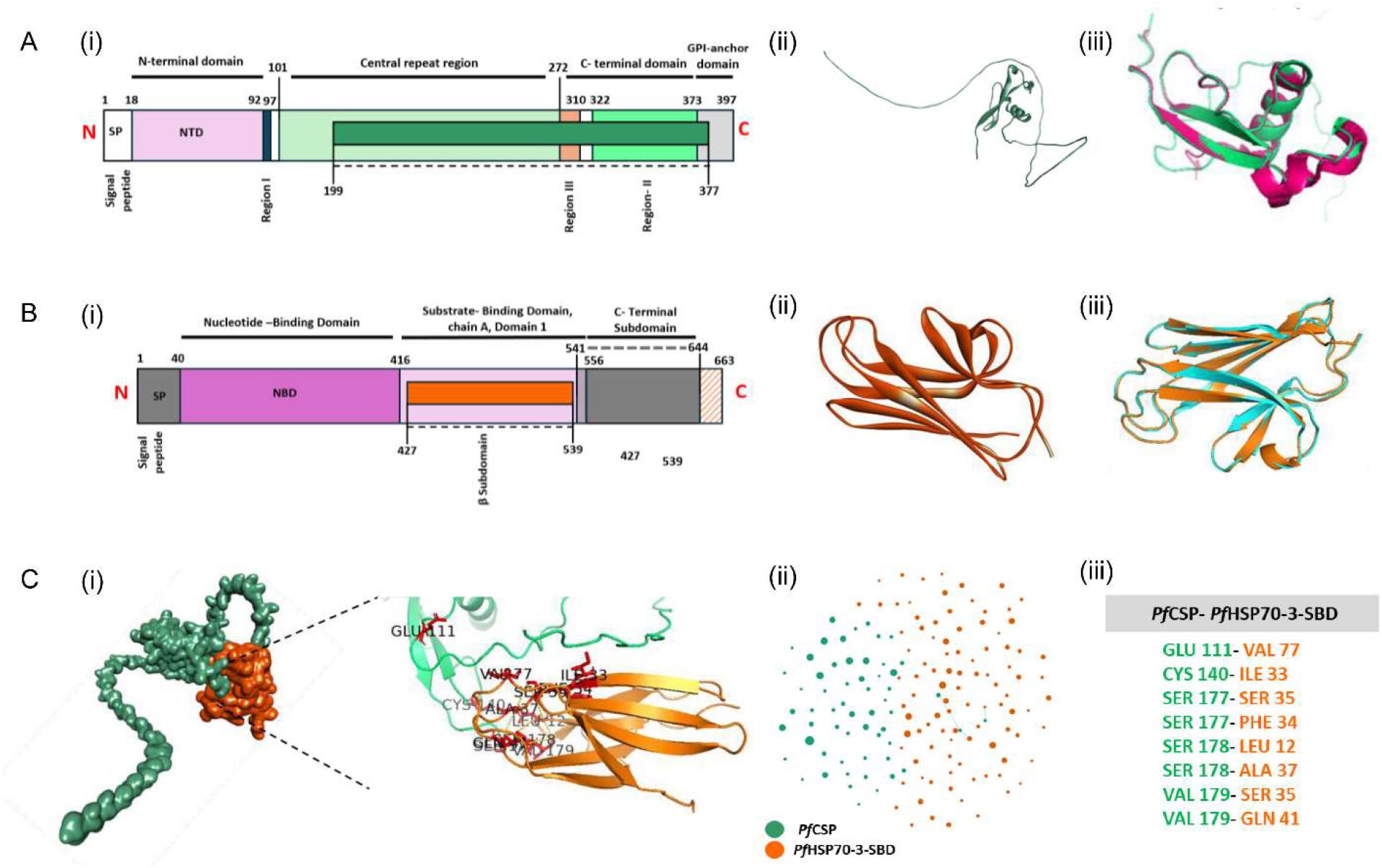
Domain organization, structural validation, and predicted *in silico* interaction between *Pf*CSP and *Pf*HSP70-3-SBD. (A) Domain architecture and structural validation of *Pf*CSP. (i) Linear domain architecture of *Pf*CSP (397 residues), comprising a signal peptide, an N-terminal domain, an intrinsically disordered central NANP/NVDP repeat region, a junction segment, and a C-terminal domain (including TSR domain), with domain boundaries are marked with residue numbers. The cloned part used in this study is highlighted by lime green box with boundary residue numbers marked on the start and end of this region connected by dashed lines. (ii) Predicted three dimensional structure of *Pf*CSP region by AlphaFold3. (iii) Structural validation via superposition of the AlphaFold3-predicted *Pf*CSP -TSR domain (lime green) onto the crystal structure of the isolated domain (PDB ID: 3VDJ, Pink) yielding a Cα RMSD of 0.315 Å. (B) Domain architecture and structural validation of *Pf*HSP70-3-SBD. (i) Linear domain architecture of *Pf*HSP70-3 (663 residues). *Pf*HSP70-3 follows the canonical HSP70 architecture of a mitochondrial transit peptide, a nucleotide-binding domain, an interdomain linker, and a C-terminal substrate-binding domain comprising the SBD β-Subdomain and a C-terminal Subdomain. The cloned SBD β-Subdomain (*Pf*HSP70-3-SBD) region used in this study is highlighted by an orange box, with boundary residue numbers are marked at the start and end of this region, connected by dashed lines. (ii) Predicted three-dimensional structure of *Pf*HSP70-3-SBD by AlphaFold3. (iii) Structural validation via superposition of the AlphaFold3-predicted *Pf*HSP70-3-SBD model (orange) onto the substrate-binding domain of the orthologous chaperone *Pf*HSP70-x (PDB ID: 6ZHI, cyan), yielding a Cα RMSD of 0.303 Å, confirming that both modelled domains closely reproduce their respective experimentally determined folds. (C) *In silico* docking and interaction analysis of the *Pf*CSP-*Pf*HSP70-3-SBD complex. (i) Cartoon representation of the docked complex using HADDOCK 2.4, with *Pf*CSP shown in lime green and *Pf*HSP70-3-SBD in orange; interface residues are shown as sticks, coloured by chain, rendered in PyMOL, illustrating the predicted binding interface. (ii) Two-dimensional interaction diagram summarizing residue-level contacts at the docked interface, derived from RING analysis. Nodes represent interface residues on *Pf*CSP (lime green) and *Pf*HSP70-3-SBD (orange). Connecting lines indicate bonds between interacting residues. (iii) List of interacting amino acid Residues for *Pf*CSP-*Pf*HSP70-3-SBD complex.

Similarly, the domain architecture for *Pf*HSP70-3 (PlasmoDB ID: *PF*3D7_1134000), defining the span of domains in the protein along with its substrate-binding domain β-subdomain, is represented by orange box (**Fig. 1B (i))**. The structural model of the substrate-binding domain β-subdomain of *Pf*HSP70-3 was similarly generated using AlphaFold3 (**Fig. 1B (ii))** and assessed using ERRAT and PROCHECK following 3Drefine refinement; ERRAT showed an overall quality factor of 93.1%, with no residue lying in the Ramachandran unfavourable region **(Fig. S2)**. Superposition of this model onto the experimentally solved substrate-binding domain of the orthologous chaperone *Pf*HSP70-x (PDB ID: 6ZHI) yielded a Cα RMSD of 0.303 Å **(Fig .1B (iii))**, further supporting the structural reliability of the *Pf*HSP70-3-SBD model.

Blind docking of the central NANP and C-terminal regions of *P. falciparum* CSP protein (*Pf*CSP) with Substrate-Binding Domain β-subdomain of *P. falciparum* HSP70-3 (*Pf*HSP70-3-SBD) in HDOCK consistently and reproducibly engaged this same folded region of *Pf*CSP as the crystal structure of *Pf*CSP αTSR domain (PDB ID: 3VDJ) across all ten output poses, independently pointing to the same interface without any restraint guidance. Restraint-driven docking in HADDOCK, guided by the AlphaFold3 interface, converged on a well-defined cluster (HADDOCK score −249.8 ± 23.0 kJ/mol; Z-score −2.3, the best among all clusters generated) that reproduced this interface with high consistency across all cluster members, engaging the same *Pf*CSP and *Pf*HSP70-3-SBD regions identified above (**Fig. 1C (i)).** The complex buried a substantial interface area (1745.5 ± 89.0 Å^2^), and the binding energetics were clearly electrostatically driven, with electrostatic contributions (−826.8 ± 91.2 kJ/mol) far exceeding van der Waals packing (−202.5 ± 13.4 kJ/mol) and a moderate desolvation penalty (78.2 ± 15.5 kJ/mol), a pattern consistent with the charged and polar character of the interface. Binding affinity, estimated using PRODIGY on the HADDOCK model, was −51.5 kJ/mol (predicted K_d_ ≈ 1 × 10⁻⁹ M), indicating a high-affinity interaction. The interface composition (22 charged-apolar, 19 polar-apolar, 3 charged-charged, and 3 polar-polar contacts) reinforces the electrostatically dominated character of the binding energetics obtained from HADDOCK. Hydrogen bonds and van der Waals contacts between interface residues of *Pf*CSP and *Pf*HSP70-3-SBD at the docked interface were visualized as a two-dimensional interaction diagram generated from RING 2.0 analysis (**Fig. 1C (ii))**, alongside the complete list of interacting amino acid residues identified for the complex (**Fig. 1C (iii))**.

To test whether this interface holds up against experimental (rather than predicted) structural data, the same docking protocols were repeated using the crystallographically solved *Pf*CSP αTSR domain (PDB ID: 3VDJ) against the AlphaFold3-predicted *Pf*HSP70-3-SBD model, in place of the AlphaFold3-predicted *Pf*CSP model used above. Blind HDOCK docking of 3VDJ against *Pf*HSP70-3-SBD converged on the same *Pf*CSP interface in all ten poses regardless of ranking, and restraint-driven HADDOCK docking of the same crystal structure against *Pf*HSP70-3-SBD reproduced this interface with high fidelity, after excluding two apparent contacts traced to a histidine-tag artifact in the crystal coordinates rather than native sequence. This experimentally anchored result closely mirrors the interface obtained from every preceding prediction-based method, providing strong independent confirmation that the interaction is not an artifact of AlphaFold3 modelling. The predicted interface is hydrogen bond-rich and predominantly electrostatic. In plain terms, the *Pf*HSP70-3-SBD (carrier protein) is predicted to grip the *Pf*CSP antigen tightly at its folded C-terminal domain.

### *In silico* docking predicts an interaction of the *Pf*CSP-*Pf*HSP70-3-SBD complex with the human CD91 receptor

The preceding section showed that *Pf*HSP70-3-SBD binds the TSR domain of *Pf*CSP through a hydrogen-bond-rich, electrostatically supported interface. For *Pf*HSP70-3-SBD to act as a carrier that presents *Pf*CSP to the immune system, two conditions must hold. The complex must remain recognizable by CD91, the receptor through which APCs take up chaperone-bound antigens.^51^ Second, CD91 should not need to interact with the surface of *Pf*HSP70-3-SBD occupied by *Pf*CSP residues. We tested this by docking the *Pf*CSP-*Pf*HSP70-3-SBD complex against the two principal ligand-binding clusters of CD91, Cluster II (**Fig 2A (i))** and Cluster IV (**Fig 2A (ii))**, both of which have comparable ligand-binding properties.^52–54^ Further, we also tested the interaction of *Pf*HSP70-3-SBD alone and *Pf*CSP alone with both the *Hs*CD91 clusters as controls. Docking used HDOCK and ClusPro, and contacts in representative ternary models were classified with RING 2.0 (**Fig. 2B (i-ii))**.

**Fig. 2:**
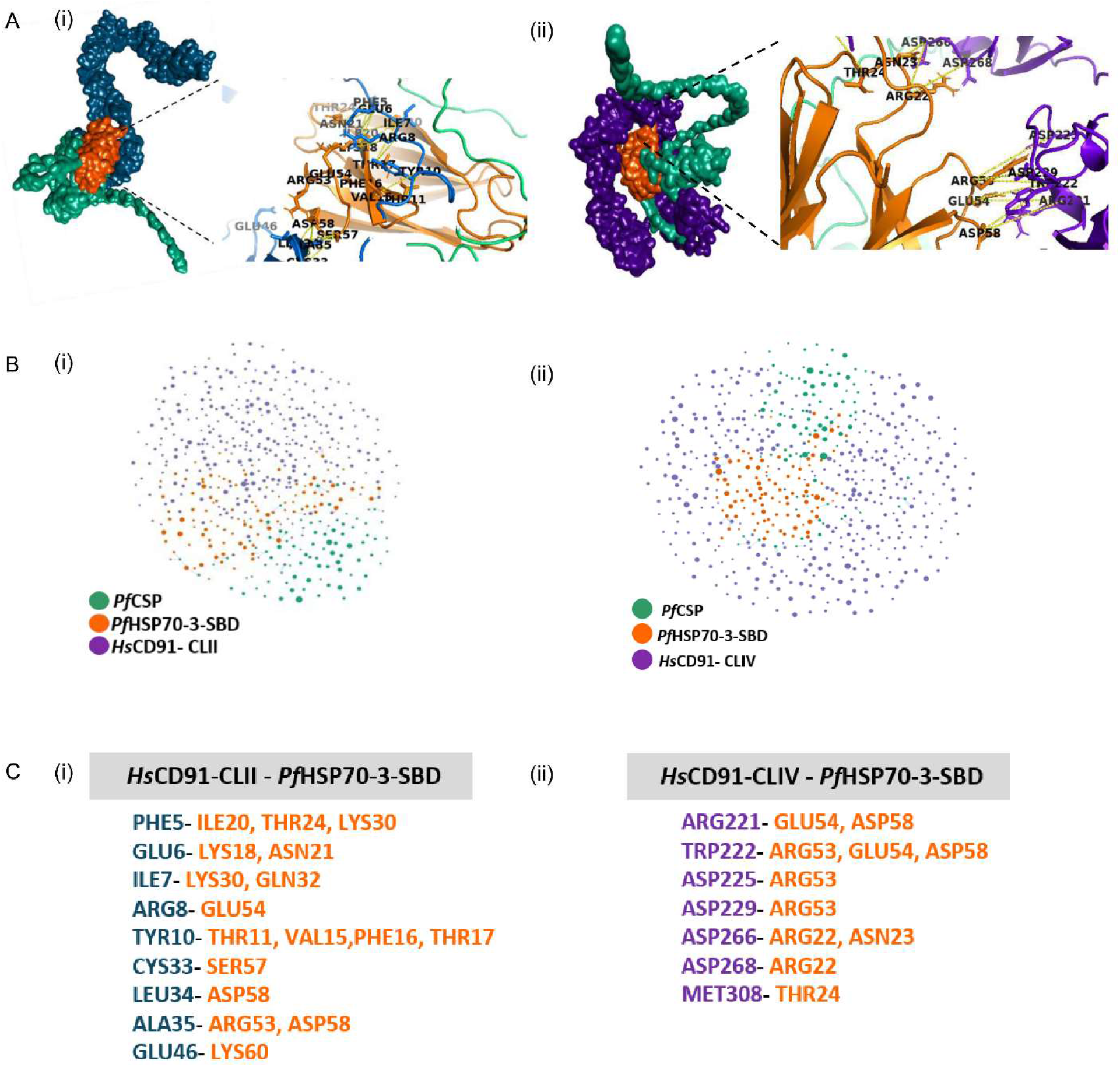
Prediction of *in silico* interaction between *Hs*CD91-CLII and *Hs*CD91-CLIV with the *Pf*CSP-*Pf*HSP70-3-SBD complex. (A) *In silico* analysis of the interaction of *Hs*CD91-CLII and HsCD91-CLIV with *Pf*CSP-*Pf*HSP70-3-SBD complex. (i) Cartoon representation of the docked complex using ClusPro, with *Hs*CD91-CLII shown in marine, *Pf*CSP in lime green, and *Pf*HSP70-3-SBD in orange; interface residues are shown as sticks, coloured by chain, rendered in PyMOL, illustrating the predicted binding interface. (ii) Cartoon representation of the docked complex using ClusPro, with *Hs*CD91-CLIV shown in purpleblue, *Pf*CSP in lime green, and *Pf*HSP70-3-SBD in orange; interface residues are shown as sticks, coloured by chain, rendered in PyMOL, illustrating the predicted binding interface. (B) 2-D Interaction Diagram for both *Hs*CD91-CLII and CLIV with *Pf*CSP-*Pf*HSP70-3-SBD complex. (i) Two-dimensional interaction diagram summarizing residue-level contacts at the docked interface, derived from RING analysis. Nodes represent interface residues on *Hs*CD91-CLII (purple) and *Pf*CSP-*Pf*HSP70-3-SBD complex [*Pf*CSP (lime green), *Pf*HSP70-3-SBD (orange)], connecting lines indicate bonds between interacting residues. (ii) Two-dimensional interaction diagram summarizing residue-level contacts at the docked interface, derived from RING analysis. Nodes represent interface residues on *Hs*CD91-CLIV (purple) and *Pf*CSP-*Pf*HSP70-3-SBD complex [*Pf*CSP (lime green), *Pf*HSP70-3-SBD (orange)], connecting lines indicate bonds between interacting residues. (C) Tabulated list for both the interactions. (i) List of interacting Amino Acid Residues for *Hs*CD91-CLII with *Pf*HSP70-3-SBD in *Pf*CSP-*Pf*HSP70-3-SBD complex. (ii) List of interacting Amino Acid Residues for *Hs*CD91-CLIV with *Pf*HSP70-3-SBD in *Pf*CSP-*Pf*HSP70-3-SBD complex.

With *Pf*HSP70-3-SBD alone, Cluster IV (HDOCK score: −212.1) scored significantly better than Cluster II (HDOCK score: −192.8). Both clusters often settled on the *Pf*CSP-binding groove, touching on average 4.7 of the 13 established *Pf*CSP-contact residues. With *Pf*CSP bound *Pf*HSP70-3-SBD, the clusters no longer differed in score (−208.7 for CLIV against −207.0 for CLII). The difference was in interference with the *Pf*CSP binding site. In all CLIV complex poses, CD91 touched at most two *Pf*HSP70-3-SBD residues that bind to *Pf*CSP, against 11 of the 13 residues across the ten poses for CLII. ClusPro also showed the same interference. We compared the poses in which CD91 buried at least 500 Å^2^ on *Pf*HSP70-3-SBD, so that low overlap was not simply a lack of contact.

We did not use buried surface area as the deciding measure, for three reasons. First, interface size is a loose guide to affinity, in a 144-complex benchmark, interface area correlated only modestly with affinity (R = 0.32).^55,56^ Second, burial was inconsistent between tools. HDOCK models of CLIV (2,742 Å^2^) buried slightly more on *Pf*HSP70-3-SBD in total than CLII (2,304 Å^2^), whereas ClusPro models of CLII (1,647 Å^2^) buried more than CLIV (681 Å^2^). Third, much of CLIV’s additional burial lay on *Pf*CSP’s NANP repeat region, which is the cargo and not the carrier. We therefore judged each cluster by the chemistry of its contacts with *Pf*HSP70-3-SBD and by its interference with the *Pf*CSP site, and treated surface area as supporting information.

The two clusters engaged *Pf*HSP70-3-SBD differently. The CLII model had a broader footprint on *Pf*HSP70-3-SBD (15 residues), but 14 of its 22 contacts were van der Waals, with 6 hydrogen bonds and 2 ionic contacts (36% charge-based). Its contacts were confined to the first 46 residues of the modelled cluster (**Fig. 2C (i))**. The Cluster IV model contacted a compact set of six *Pf*HSP70-3-SBD residues (Arg22, Asn23, Thr24, Arg53, Glu54, Asp58), all outside the *Pf*CSP-binding site. It used 8 hydrogen bonds, 3 ionic contacts (≤4.0 Å),^57^ and 1 cation–π contact, so 57% of its contacts were charge-based (**Fig. 2C (ii))**. Its CD91 contacts spread over about 210 residues spanning several repeats, in line with the multi-repeat engagement described for this receptor.^38^ Contact type describes the character and selectivity of recognition, not affinity.^55^ On that basis, CLIV recognizes *Pf*HSP70-3-SBD more selectively through charge-based contacts, whereas Cluster II makes broader, weaker-type contact closer to the *Pf*CSP-binding site.

The CD91 residues in the Cluster IV model that contact *Pf*HSP70-3-SBD (Asp3556, Asp3560, Asp3597, Asp3599) are acidic residues of the Ca²⁺-coordinating cores described for this cluster.^38^ They pair with *Pf*HSP70-3-SBD arginines in the manner of the canonical acidic-pocket mode of LDL-receptor-family members.^58^ In both clusters, CD91 contact with *Pf*CSP was limited to hydrogen bonds and van der Waals contacts, with no ionic contacts, so charge-based anchoring was directed only at *Pf*HSP70-3-SBD.

Taken together, these predictions support a division of labor in which CD91 recognizes *Pf*HSP70-3-SBD as the carrier while *Pf*CSP remains bound and makes only secondary, non-ionic contact. The primary CD91 engagement site for the *Pf*CSP-*Pf*HSP70-3-SBD complex is CLIV, while CLII acts as a secondary site. In plain terms, it appears that CD91 is able to grab the carrier (*Pf*HSP70-3-SBD) at a site away from where the antigen is held, and Cluster IV does this most selectively. However, we do not claim higher affinity than Cluster II. This is the arrangement *Pf*HSP70-3-SBD needs to deliver *Pf*CSP to CD91-bearing antigen-presenting cells, and it gives a structural basis for developing the *Pf*CSP-*Pf*HSP70-3-SBD complex as a vaccine co-formulation capable of enhancing antigen cross-presentation.

### Expression and purification of *Pf*CSP and *Pf*HSP70-3-SBD recombinant proteins

To confirm the *in vitro* interaction of *Pf*HSP70-3-SBD and *Pf*CSP, and study the adjuvant effect of *Pf*HSP70-3-SBD in amplifying the antigen-specific immune responses to *Pf*CSP antigen, recombinant proteins were purified. The recombinant expression vectors pMTSAra-*Pf*HSP70-3-SBD and pET-28b(+)-*Pf*CSP encode *Pf*HSP70-3-SBD and *Pf*CSP proteins, respectively. The recombinant proteins were hexahistidine tagged and affinity chromatography was used for their purification. The substrate-binding domain β-subdomain of *Pf*HSP70-3 was purified using *E. coli* expression system. The recombinant *Pf*HSP70-3-SBD protein, with a C-terminal His tag, was obtained as a single, homogeneous band of high purity at the expected molecular weight of 14.06 kDa on a CBB-stained SDS-PAGE gel. Western blot analysis using anti-His antibody further confirmed the identity of the recombinant *Pf*HSP70-3-SBD protein (**Fig. 3A**). We next purified the region of *Pf*CSP corresponding to the central repeat and C-terminal domains. This region was used as the pre-erythrocytic stage vaccine antigen of *P. falciparum* for recombinant protein purification. The purified recombinant *Pf*CSP, along with an N-terminal His-tag, was obtained as a single homogeneous band of high purity at a molecular weight slightly higher (∼30 kDa) than its predicted size of 22.92 kDa, as detected by CBB-stained SDS-PAGE and western blotting with an anti-His antibody (**Fig. 3B**). Both recombinant proteins were subsequently used for protein-protein interaction studies and animal immunization for assessment of antigen-specific immune responses.

**Fig. 3:**
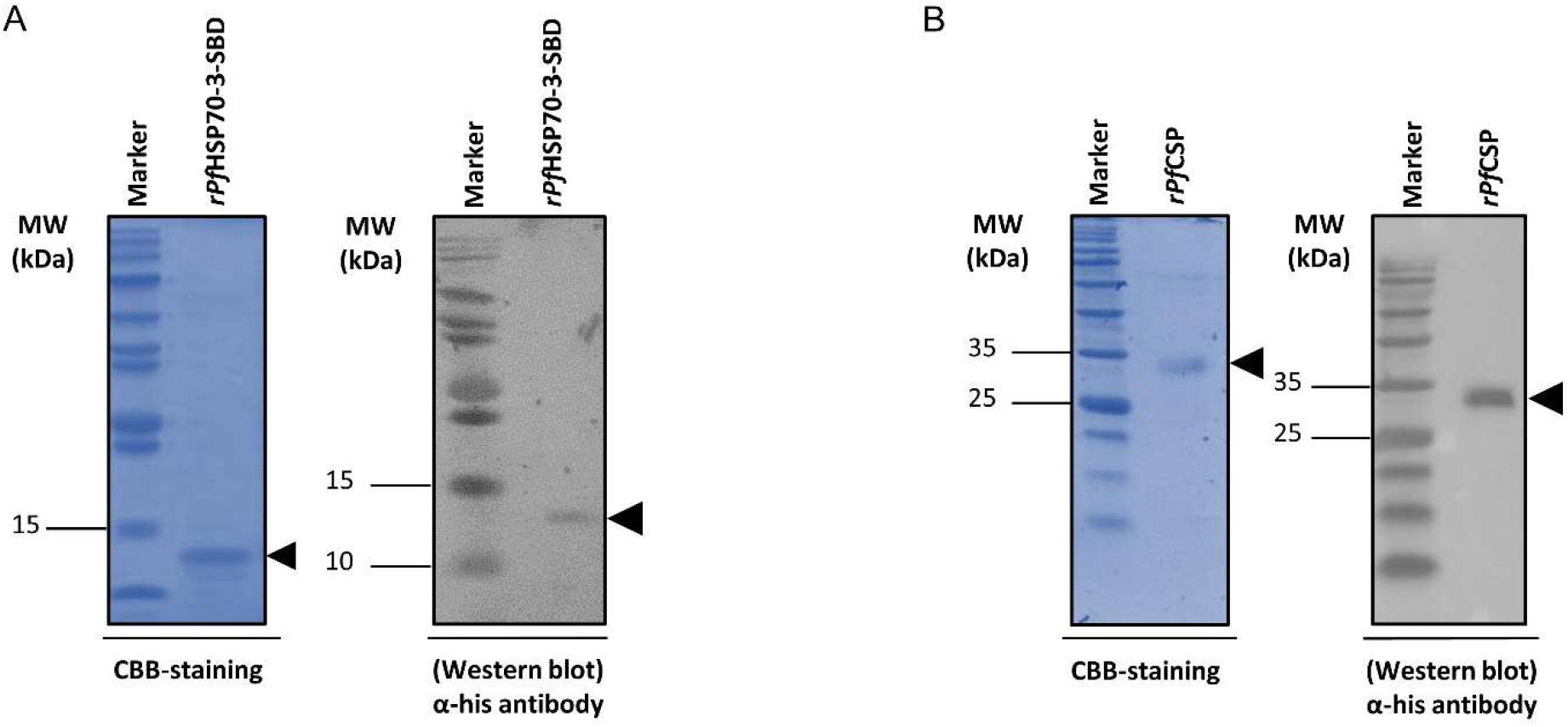
Purification and validation of recombinant *Pf*HSP70-3-SBD and *Pf*CSP proteins. (A) Purity and identity of Ni-NTA-purified hexahistidine-tagged recombinant *Pf*HSP70-3-SBD (14.06 kDa) assessed by CBB-stained SDS-PAGE and Western blotting using an anti-His antibody. (B) Purity and identity of Ni-NTA-purified hexahistidine-tagged recombinant *Pf*CSP assessed by CBB-stained SDS-PAGE and Western blotting using an anti-His antibody. *Pf*CSP migrated at an apparent molecular weight of ∼30 kDa, higher than its theoretical molecular weight of 22.92 kDa. Molecular weight (MW) markers are shown to the left in kilodaltons (kDa). Arrows indicate the bands corresponding to the molecular weights of *Pf*HSP70-3-SBD and *Pf*CSP recombinant proteins.

### *P. falciparum* CSP interacts with the substrate-binding domain of *Pf*HSP70-3

With the understanding that the substrate-binding domain of HSPs is responsible for protein binding and is fundamental for HSP70-mediated cross-presentation,^29–31,59^ we investigated whether *Pf*HSP70-3-SBD interacts with *P. falciparum* CSP. With an established *in silico* binding, we purified the recombinant proteins *Pf*HSP70-3-SBD and *Pf*CSP, and evaluated their interaction using ELISA and far-western dot blot analysis. An ELISA was designed in which 200 ng/well *Pf*HSP70-3-SBD was coated onto the ELISA plate, followed by overlay with varied concentrations of *Pf*CSP protein, and binding was detected using rabbit mAb 2A10 (**Fig. 4A**). The results indicated a strong interaction between the substrate-binding domain of *Pf*HSP70-3 and *P. falciparum* CSP. Interaction was dose-dependent, and higher concentrations of *Pf*CSP exhibited enhanced interaction with *Pf*HSP70-3-SBD. The strongest interaction corresponded to the highest mean OD_450_ value of 3.41, which was obtained with 2,000 ng of *Pf*CSP and 200 ng of *Pf*HSP70-3-SBD. On the other hand, negligible interaction was detected with the respective concentrations of BSA and *Pf*CSP (OD_450_: 0.69), which served as the negative control in the experiment (**Fig. 4B**). To further validate these findings, far-western dot blot analysis was performed and ImageJ was used to perform the densitometry analysis. Blot intensities were measured quantitatively and arbitrary densitometry units (ADUs) were plotted in GraphPad Prism 8.0. *Pf*HSP70-3-SBD (spotted) produced a prominent signal corresponding to the interaction with *P. falciparum* CSP (overlaid) (ADU: 343.43), whereas no interaction was detected between BSA and *Pf*CSP, which was used as the negative control (ADU: 1.00) (**Fig. 4C (i))**. Similarly, *Pf*CSP (spotted) also produced a prominent signal upon interaction with *Pf*HSP70-3-SBD (overlaid), with a higher ADU of 7.60, as compared to the interaction between BSA and *Pf*HSP70-3-SBD used as the negative control (ADU: 1.00) (**Fig. 4C (ii))**. With these results, we hypothesize that *Pf*CSP antigen potentially binds to *Pf*HSP70-3-SBD, which may facilitate an efficient cross-presentation of the bound antigen, thereby augmenting antigen-specific immune responses. Collectively, these findings support a direct or indirect interaction between *Pf*HSP70-3-SBD and the *P. falciparum* antigen CSP, providing a potential molecular basis for the enhanced immunogenicity observed following a co-formulation.

**Fig. 4:**
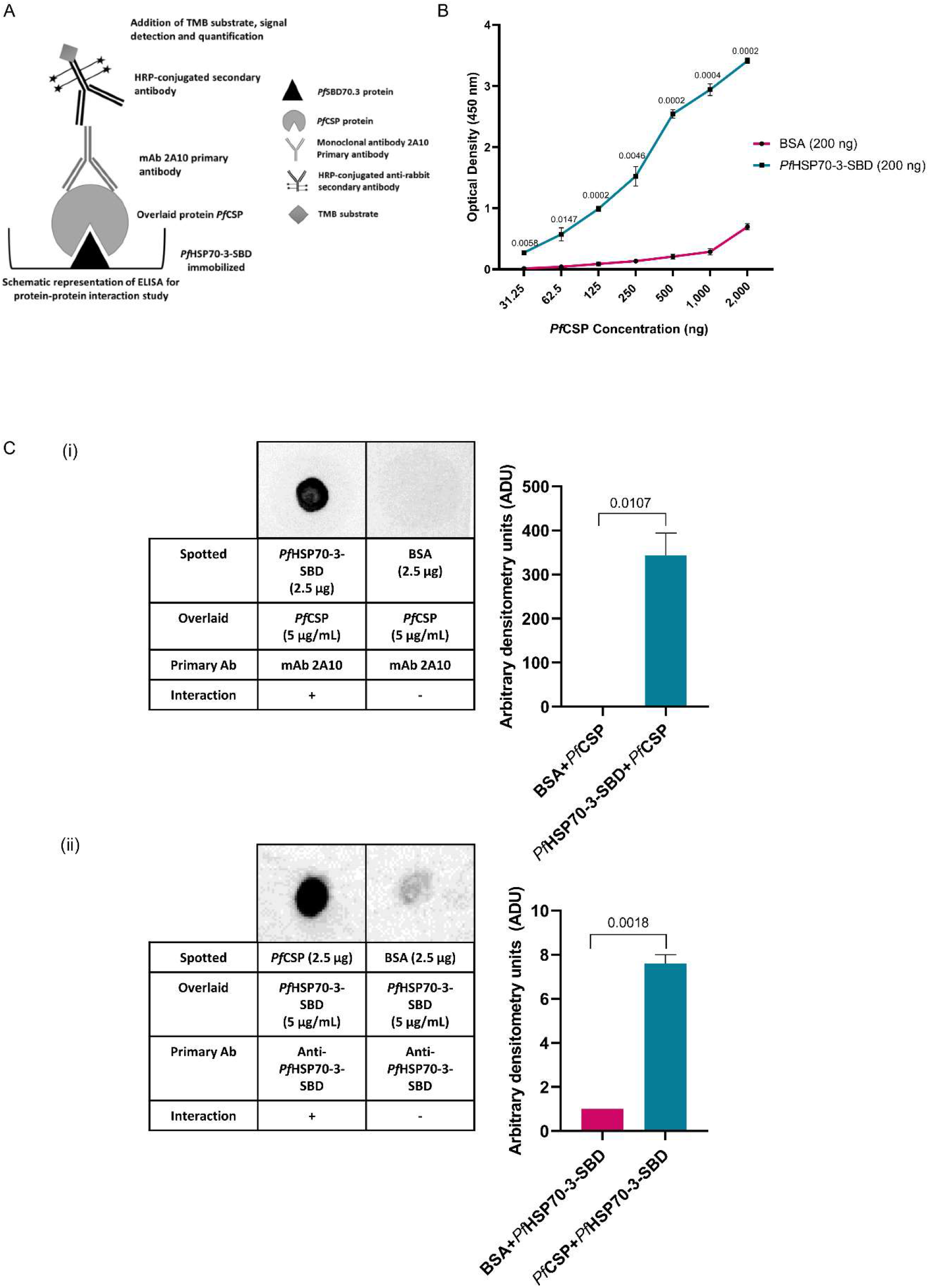
*In vitro* biochemical characterization of *Pf*HSP70-3-SBD and *Pf*CSP interaction. (A) Schematic representation of ELISA designed to assess the interaction of *Pf*HSP70-3-SBD and *Pf*CSP. (B) The *in vitro* interaction between the SBD of *P. falciparum* HSP70-3 and *P. falciparum* CSP, confirmed by ELISA. 200 ng *Pf*HSP70-3-SBD recombinant protein was immobilized per well, and *Pf*CSP recombinant protein was overlaid at different concentrations (31.25 ng, 62.50 ng, 125 ng, 250 ng, 500 ng, 1,000 ng, 2,000 ng). BSA (as an overlaid protein) served as the negative control. The mean ± standard deviation (SD) from triplicate experiments was used to represent the data. Statistical analysis was performed by comparing the respective concentrations of experimental groups with the BSA control using Student’s unpaired two-tailed t-test. (C) Representative images and analysis of Far-Western dot blot for the interaction between *Pf*HSP70-3-SBD and *Pf*CSP. Reciprocal binding assays were performed using (i) *Pf*HSP70-3-SBD (spotted protein) and *Pf*CSP (overlaid protein), and (ii) *Pf*CSP (spotted protein) and *Pf*HSP70-3-SBD (overlaid protein). BSA was immobilized and served as a negative control during the experiments. ImageJ software was used to quantify blot intensities, and arbitrary densitometry units (ADUs) were calculated and plotted using GraphPad Prism 8. Student’s unpaired two-tailed t-test was used for statistical analysis by comparing the experimental groups with the BSA control. p-values are displayed, where p < 0.05 was considered significant and p-values > 0.05 were considered statistically insignificant.

### Enhancement of humoral immune responses to the malaria liver-stage antigen *Pf*CSP with *Pf*HSP70-3-SBD co-formulation

Since the adjuvant-like potential of a *Pf*HSP70 fragment-based DNA construct has already been demonstrated in an earlier study,^26^ we examined the effect of the substrate-binding domain of *Pf*HSP70-3 as a protein-based adjuvant in amplifying the antigen-specific antibody responses when co-formulated with the *P. falciparum* CSP antigen. BALB/c mice were immunized intramuscularly with *Pf*CSP antigen formulated in alum, either with or without *Pf*HSP70-3-SBD (**Fig. 5A**). One prime and two booster doses were administered at two-week intervals. As a control, mice were immunized with PBS formulated in alum. Blood was collected after each immunization at designated time points, and sera were isolated to evaluate the anti-*Pf*CSP total IgG responses. To investigate the effectiveness of *Pf*HSP70-3-SBD in enhancing the immune response specific to *Pf*CSP, indirect ELISA was performed. Minimal or no detectable *Pf*CSP-specific IgG responses were observed in mice immunized with PBS/Alum alone on days 13, 27 and 42. Furthermore, *Pf*CSP/Alum in combination with *Pf*HSP70-3-SBD (*Pf*CSP/*Pf*HSP70-3-SBD/Alum) was found to elicit enhanced anti-*Pf*CSP IgG titers compared to the *Pf*CSP/Alum immunized group. The enhancement of anti-*Pf*CSP IgG titers upon co-formulation of *Pf*HSP70-3-SBD was consistently observed after every immunization. Following the second booster dose of *Pf*CSP/*Pf*HSP70-3-SBD/Alum, peak anti-*Pf*CSP total IgG levels were detected. Notably, the *Pf*CSP/*Pf*HSP70-3-SBD/Alum-immunized group exhibited significantly higher mean OD_450_ values across all serum dilutions on days 27 and 42, when compared with the *Pf*CSP/Alum-immunized group (**Fig. 5B**). Although adsorption of protein subunit antigens with alum is efficient in inducing humoral immune responses, these findings further support the additive role of *Pf*HSP70-3-SBD in augmenting *Pf*CSP-specific humoral immunity, which may contribute to strong protective immunity against the liver stage of the *P. falciparum* life cycle.

**Fig. 5:**
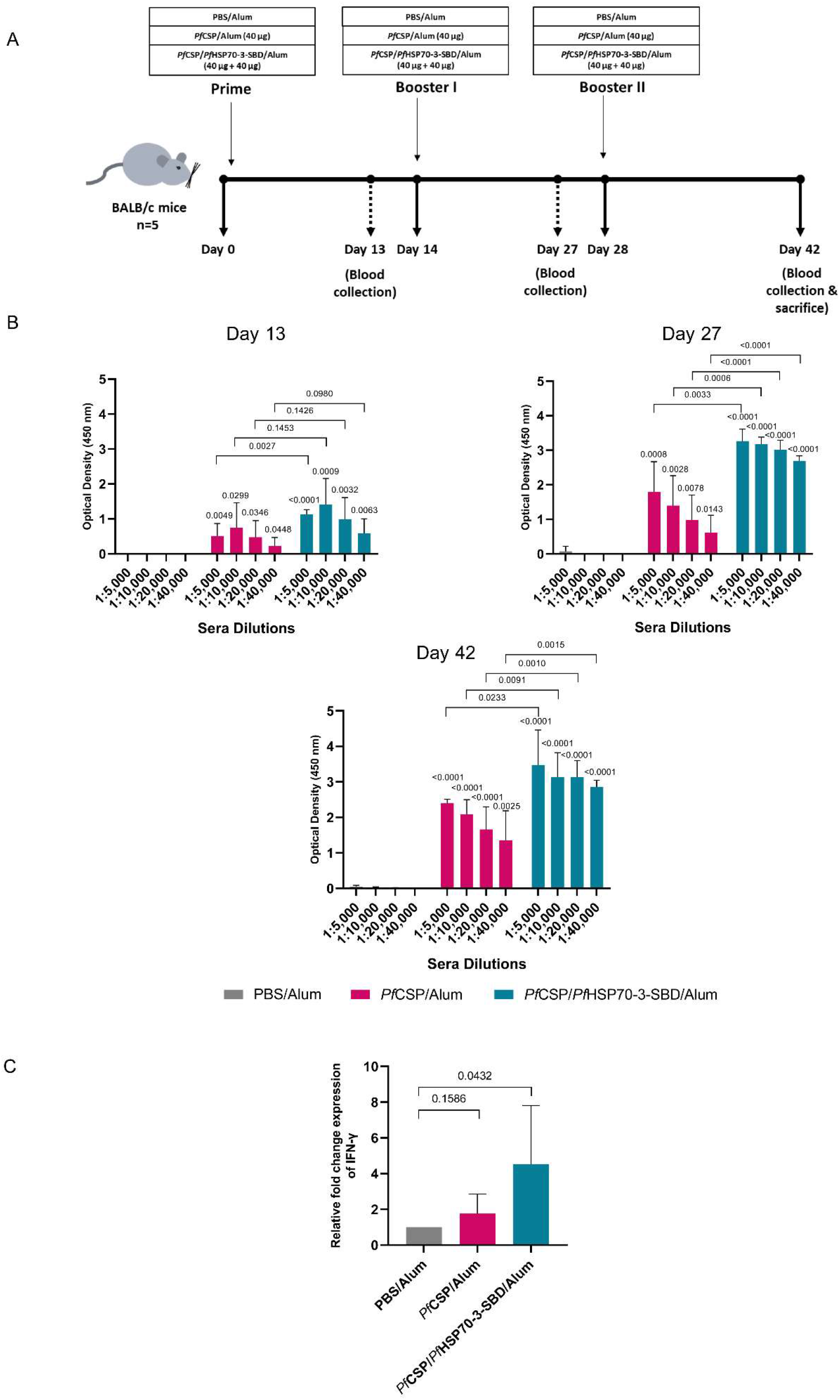
Adjuvant effect of *Pf*HSP70-3-SBD co-formulated with *Pf*CSP antigen in enhancing humoral and cellular immune responses. (A) Scheme for animal immunization showing the experimental groups, dosage, and timeline. *Pf*CSP/Alum or *Pf*CSP/*Pf*HSP70-3-SBD/Alum were administered intramuscularly to BALB/c mice (n=5 per group). Control mice received PBS/Alum only. Mice were immunized with one prime and two booster doses at two-week intervals. Blood collection was done on days 13, 27, and 42 to isolate respective serum samples. (B) Induction of total anti-*Pf*CSP IgG responses in BALB/c mice following each immunization. Total IgG antibody responses against *Pf*CSP antigen were evaluated by indirect ELISA using 200 ng of *Pf*CSP protein per well for coating. Sera collected from five mice per group on days 13, 27, and 42 were pooled separately for each time point. Serum samples were diluted serially (1:500 to 1:40,000), and OD_450_ values were plotted using GraphPad Prism version 8.0 software. The experiment was performed twice in triplicate, and data are presented as mean ± standard deviation (SD). Statistical comparisons were performed using an unpaired two-tailed Student’s t-test, comparing each experimental group with the PBS/Alum control group and the *Pf*CSP/Alum group with the *Pf*CSP/*Pf*HSP70-3-SBD/Alum group. (C) IFN-γ mRNA expression in mouse splenic tissues following immunization with *Pf*CSP/*Pf*HSP70-3-SBD/Alum. On day 42, mice were sacrificed, and RNA was isolated from whole spleens. mRNA expression was quantified from splenic tissues using qRT-PCR. Expression levels of the IFN-γ cytokine were normalized using the GAPDH housekeeping gene and quantified using the 2^-ΔΔCt^ method. Data are represented as mean ± standard deviation (SD) and plotted using GraphPad Prism version 8.0 software. For statistical analysis, the experimental groups were compared with the PBS/Alum control group, and Student’s unpaired two-tailed t-test was used. p-values < 0.05 were considered statistically significant, and p > 0.05 were considered insignificant.

### Co-formulation of *Pf*HSP70-3-SBD with the *Pf*CSP antigen enhances IFN-γ expression in mice

Following the assessment of the humoral responses induced by immunization with *Pf*CSP/*Pf*HSP70-3-SBD/Alum, we determined whether the incorporation of *Pf*HSP70-3-SBD into the antigen formulation enhanced cellular immune responses. We measured the relative fold-change in the expression of IFN-γ, a pro-inflammatory cytokine, using splenic tissues collected from control and immunized mice 14 days after the second booster dose. The fold-change was calculated relative to the PBS/Alum-immunized group (set at 1.00) and normalized to GAPDH. We found that the *Pf*CSP/Alum-immunized group exhibited increased IFN-γ expression (1.76-fold) compared with the PBS/Alum-immunized group. Interestingly, immunization with *Pf*CSP/*Pf*HSP70-3-SBD/Alum further resulted in a significantly higher expression of IFN-γ (4.52-fold) (**Fig. 5C**). These findings demonstrate that incorporation of *Pf*HSP70-3-SBD into the *Pf*CSP/Alum formulation enhances IFN-γ expression, suggesting that *Pf*HSP70-3-SBD promotes a stronger Th1-associated immune response which is associated with effective parasite control and clearance.

### Recombinant antisera specific to *Pf*CSP/*Pf*HSP70-3-SBD/Alum reacts with *Pf*CSP and neutralizes *Pf/Pb* CSP transgenic sporozoites

To demonstrate the specificity of *Pf*CSP/*Pf*HSP70-3-SBD/Alum antisera to *Pf*CSP repeats, lysates were generated from *Pf/Pb* CSP transgenic sporozoites and western blotting was performed. The immune sera identified a distinct band of ∼54 kDa corresponding to *Pf*CSP in the transgenic sporozoites (**Fig. 6A**). This implied that antisera against *Pf*CSP/*Pf*HSP70-3-SBD/Alum retained it’s the ability to recognize native *Pf*CSP repeats expressed by the transgenic sporozites.

**Fig. 6:**
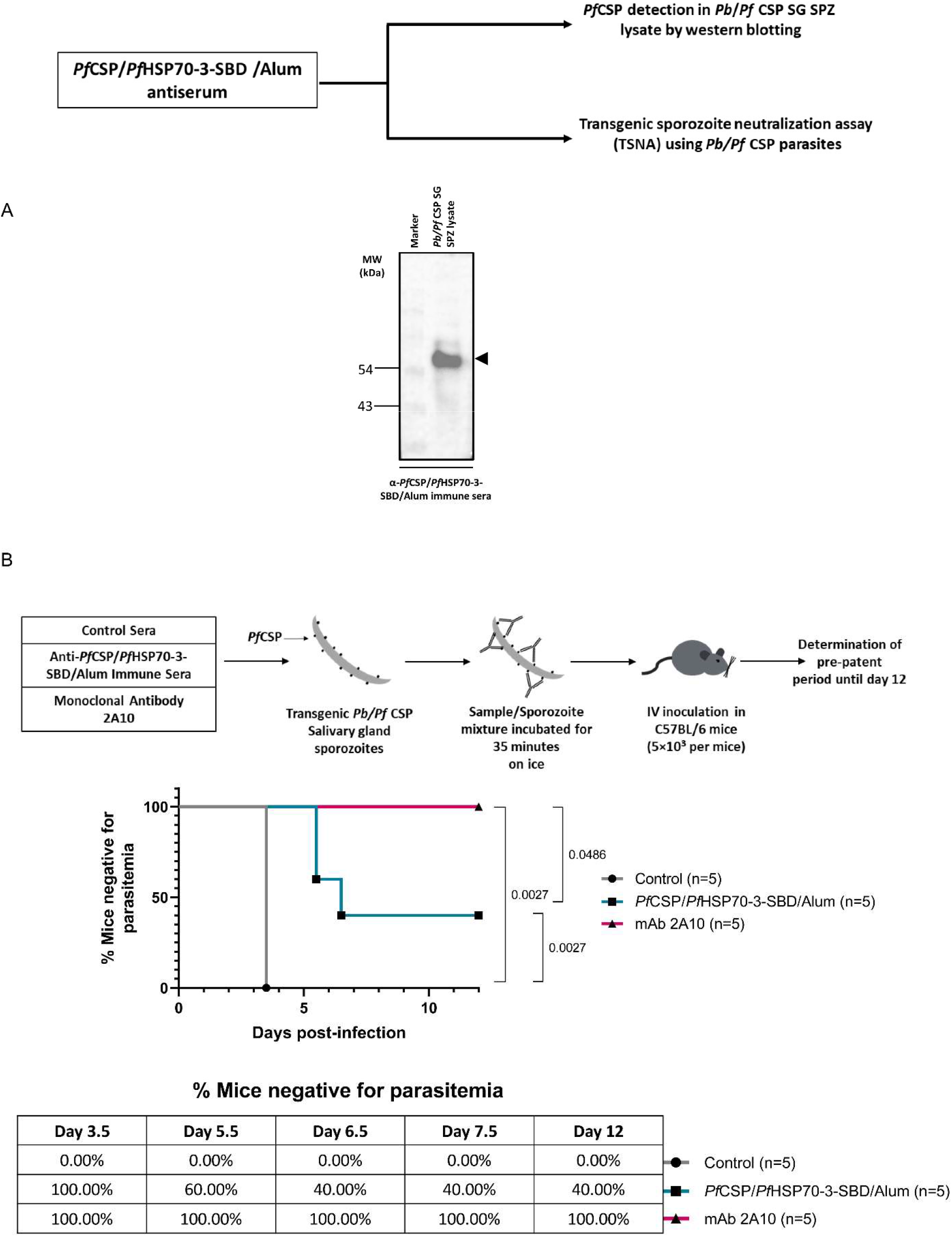
Functional evaluation of anti-*Pf*CSP/*Pf*HSP70-3-SBD/Alum immune sera using transgenic *Pb/Pf* CSP sporozoites. (A) Representative blot showing recognition of *Pf*CSP antigen by anti-*Pf*CSP/*Pf*HSP70-3-SBD/Alum immune sera in transgenic *Pb/Pf* CSP sporozoite lysates. Immune sera from *Pf*CSP/*Pf*HSP70-3-SBD/Alum-immunized mice collected on day 42 post-immunization were evaluated for recognition of parasite-expressed *Pf*CSP antigen. Lysates of *Pb/Pf* CSP sporozoites were prepared and analyzed by Western blotting. Molecular weight markers are indicated on the left in kilodaltons (kDa). The black arrow indicates the band corresponding to *Pf*CSP (∼54 kDa). (B) Transgenic sporozoite neutralization assay (TSNA) using anti-*Pf*CSP/*Pf*HSP70-3-SBD/Alum immune sera. Sera from BALB/c mice (n = 5) immunized with one prime and two booster doses of *Pf*CSP/*Pf*HSP70-3-SBD/Alum were pooled and evaluated by TSNA. *Pb/Pf* CSP salivary gland sporozoites were pre-incubated with anti-*Pf*CSP/*Pf*HSP70-3-SBD/Alum immune sera (closed square) or control sera (closed circle) at 1:10 dilution. Sera/sporozoite mixtures were inoculated intravenously into naïve C57BL/6 mice (n=5 per group). The pre-patent period was monitored up to day 12 by examining Giemsa-stained thin blood smears under a microscope. The delay in the pre-patent period was compared relative to the control group. Sporozoites pre-incubated with the *Pf*CSP repeat-specific monoclonal antibody 2A10 (closed triangle) were included as a positive control (1:50 dilution of 1 mg/mL concentration). Percentage of mice negative for the pre-patent parasite is shown in the table. Kaplan-Meier curves were generated using GraphPad Prism 8.0 to show the percentage of mice remaining negative for blood-stage parasites over time. Statistical analysis was performed by the Log-rank (Mantel-Cox) test, and p-values were shown with respect to the control group. p-values are indicated, with p < 0.05 considered statistically significant.

We further evaluated the sporozoite-neutralizing activity of *Pf*CSP/*Pf*HSP70-3-SBD/Alum specific sera using transgenic *Pb/Pf* CSP parasites. A transgenic sporozoite neutralization assay (TSNA) was set up to assess the ability of *Pf*CSP/*Pf*HSP70-3-SBD/Alum induced antibodies to inhibit sporozoite infectivity. In this assay, *Pb/Pf* CSP transgenic sporozoites were pre-incubated with sera from *Pf*CSP/*Pf*HSP70-3-SBD/Alum-immunized mice and inoculated intravenously into naïve C57BL/6 mice, 35 minutes post-neutralization. Controls included exposure of *Pf/Pb* sporozoites to either non-immune sera or 2A10. Giemsa-stained blood smears were examined daily from all mice exposed to treated sporozoites, from day 3 post-infection, to monitor blood-stage parasitemia. While all mice infected with sporozoites treated with non-immune sera developed pre-patency by day 3.5, no breakthrough was detected in any mice receiving the *Pf*CSP/*Pf*HSP70-3-SBD/Alum immune sera/sporozoite mixture until day 4 post-infection. However, on day 5.5, 60.00% of these mice remained negative for blood-stage parasites, and the efficacy decreased to 40.00% by day 6.5 and day 12 (**Fig. 6B, Table S3)**. The prepatent period was longer in the *Pf*CSP/*Pf*HSP70-3-SBD/Alum group than in the control group. The C57BL/6 mice inoculated with sporozoites incubated with control/non-immune sera developed blood-stage parasitemia by day 3.5. The *Pf*CSP repeat-specific monoclonal antibody 2A10, used as a positive control, completely prevented breakthrough blood-stage parasitemia in all mice until day 12. A delay in the pre-patent period was interpreted as evidence of sporozoite-neutralizing activity by vaccine-induced antibodies. The delay in the prepatent period observed with *Pf*CSP/*Pf*HSP70-3-SBD/Alum immune sera, as compared with unimmunized control sera, indicated a modest neutralization potential. Long-term monitoring revealed 40.00% survival in the *Pf*CSP/*Pf*HSP70-3-SBD/Alum group, compared with 100% survival in the mAb 2A10 (positive control) group at 2 months post-infection with no detectable parasitemia. These results point to a partially efficacious nature of *Pf*CSP/*Pf*HSP70-3-SBD/Alum in inducing immunity against the sporozoite stages.

## 4. DISCUSSION

The development of a highly effective vaccine against any infectious disease is a cornerstone of public health policy and remains exceptionally challenging. To prevent the clinical manifestations of infection, an ideal vaccine requires the induction of both B cell- and T cell-dependent immune mechanisms without the risk of anaphylaxis or immune exhaustion.^60^ Vaccination with an optimized formulation of antigens and adjuvants provides durable and efficacious protection from disease. Patterns or components common on pathogen surfaces led to the discovery of novel adjuvants based on pathogen-associated molecular patterns (PAMPs). These patterns are recognized by pattern recognition receptors (PRRs) on the surface of immune cells, activating the innate immune responses that subsequently direct adaptive immunity.^61^ A number of ligands that target PRRs (e.g., toll-like receptors) have been extensively characterized as vaccine adjuvants.^62–65^ There remains strong demand for pathogen-derived vaccine adjuvants to elicit robust immune responses against intracellular pathogens, with several undergoing clinical trials.^66^

For prevention of infectious diseases that require the induction of a cellular immune response, molecular chaperones have been proposed as a new class of adjuvants in the design of vaccine formulations. Although chemical chaperones, including 4-phenylbutyric acid (PBA) and tauroursodeoxycholic acid (TUDCA) for type 2 diabetes treatment and green tea catechins for cancer prevention, have shown therapeutic potential, they are limited by several intrinsic drawbacks, including toxicity, off-target interactions, and pharmacokinetic inefficiency.^67–69^ Unlike chemical chaperones, molecular chaperones, including HSPs, serve as a quality-control system in addition to the protein-folding process.^70^ Besides maintaining protein homeostasis, the unique capacity of HSPs to shuttle proteins/peptides and facilitate cross-presentation of antigens has allowed their use in multiple vaccine applications. Professional antigen-presenting cells (APCs), especially dendritic cells, can effectively internalize HSP-associated antigens primarily via CD91 receptor-mediated mechanisms.^51^ This interaction can promote APC activation and functional maturation, thereby enhancing their capacity to prime naïve T cells. In general, exogenous antigens are conventionally processed through the MHC class II pathway of APCs, leading primarily to the activation of antigen-specific CD4⁺ T lymphocytes. However, the uptake of HSP-peptide complexes by professional APCs facilitates intracellular routing of exogenous antigens toward the MHC class I presentation pathway, supporting cross-presentation and subsequent activation of antigen-specific CD8⁺ T-cell responses. This mechanism may promote the activation of antigen-specific CD8⁺ T cells in addition to CD4⁺ T-cell responses. This provides a mechanistic basis for the potential of HSPs to induce coordinated cellular and humoral immune responses and supports their use as adjuvanting components in vaccine formulations.^21,71^

Although a previous study has reported lack of immunomodulatory activity of *Plasmodium falciparum* HSP70s,^72^ we strategically explored the SBD β-subdomain (peptide-binding region) of *Pf*HSP70-3 as a functional module for antigen delivery. We hypothesized that its peptide-binding capacity could facilitate the uptake and intracellular processing of associated antigenic peptides, thereby promoting their cross-presentation through the MHC-I pathway and enhancing antigen-specific CD8⁺ T-cell responses. In this regard, we explored the substrate-binding domain of *P. falciparum* HSP70-3 as a protein-based adjuvant. To date, the circumsporozoite protein of *P. falciparum* has represented a significant milestone as a pre-erythrocytic stage antigen in malaria control through RTS,S/AS01 and R21/Matrix-M vaccines recommended by WHO.^4,5^ We therefore co-formulated *Pf*HSP70-3-SBD with the *Pf*CSP antigen to investigate its potential to enhance *Pf*CSP-specific immune responses.

We first determined the *Pf*CSP antigen-binding capacity of the substrate-binding domain β-subdomain of *Pf*HSP70-3. An *in silico* protein-protein docking analysis provided a plausible molecular basis for the interaction between *Pf*HSP70-3-SBD and *Pf*CSP. Docking analysis revealed that the TSR domain of *Pf*CSP interacts with a binding pocket within the β-subdomain of *Pf*HSP70-3 substrate-binding domain. The predominance of hydrophobic residues at the predicted interface of *Pf*HSP70-3-SBD, together with contributions from Ser35 and Gln41, may facilitate the association and stabilization of the *Pf*CSP-*Pf*HSP70-3-SBD complex. This was further supported by *in vitro* biochemical characterization, where we demonstrated a specific interaction between recombinant *Pf*CSP and *Pf*HSP70-3-SBD proteins, as evidenced by ELISA and Far-Western dot blot assays. Although majority of ligands are known to interact predominantly with clusters II and IV of host CD91,^52^ our *in silico* analyses predicted interactions of the *Pf*CSP-*Pf*HSP70-3-SBD complex with both receptor clusters, with a primary engagement with cluster IV in the ternary complex. Notably, docking of *Pf*CSP alone with either cluster did not yield a docking output under the conditions examined. This differential docking behaviour raises the possibility that *Pf*HSP70-3-SBD-bound *Pf*CSP antigen may engage CD91 receptor on the surface of APCs, potentially providing a mechanistic link to receptor-mediated antigen uptake. Such an interaction could be relevant to the intracellular routing of exogenous antigen for cross-presentation, thereby offering a plausible explanation for the antigen-specific immune responses observed following co-formulation of *Pf*HSP70-3-SBD with *Pf*CSP. Nevertheless, the findings are computational and docking was rigid-body on modeled receptor fragments. This does not establish a direct receptor engagement or demonstrate CD91-dependent internalization and cross-presentation; therefore, targeted experimental studies are required to validate this proposed mechanism. Thus, this chaperonic domain-specific adjuvant strategy offers potential advantages, including (i) an antigen-delivery function; (ii) the potential to enhance antigen cross-presentation; and (iii) broad applicability owing to its potential for co-formulation with a wide range of vaccine antigens.

Since alum is well known to augment humoral immunity, we investigated the adjuvanting properties of *Pf*HSP70-3-SBD in combination with alum to determine whether their combination could further enhance antibody- and cell-mediated immune responses. In the present study, co-formulation of *Pf*CSP/Alum with *Pf*HSP70-3-SBD elicited enhanced humoral, characterized by *Pf*CSP-specific total IgG, and cellular, characterized by IFN-γ overexpression, immune responses compared with the formulation lacking *Pf*HSP70-3-SBD. The enhanced IFN-γ response observed following co-formulation with *Pf*HSP70-3-SBD is particularly relevant, as IFN-γ is a pro-inflammatory cytokine that plays a critical role in anti-sporozoite immunity and the control of liver-stage malaria infection.^73^ We also performed western blot analysis using lysates of transgenic *Pb/Pf* CSP sporozoites and demonstrated that immune sera from mice immunized with the *Pf*CSP/*Pf*HSP70-3-SBD/Alum formulation efficiently recognized parasite-expressed *Pf*CSP. In co-formulated protein subunit vaccines, interactions between a protein-based adjuvant and antigen may influence the structural conformation and accessibility of the neutralizing epitopes of antigen. Such interactions could potentially alter the exposure of immunologically relevant determinants and thereby influence B-cell recognition and the magnitude or specificity of antigen-specific antibody responses. In this regard, we performed a transgenic sporozoite neutralization assay using immune sera and *Pb/Pf* CSP sporozoites. The ability of immune sera to neutralize transgenic *Pb/Pf* CSP sporozoites and prolong the pre-patent period indicates that co-formulation and interaction between *Pf*HSP70-3-SBD and *Pf*CSP appear to retain exposure of *Pf*CSP neutralizing epitopes. This further supports the functional relevance of *Pf*HSP70-3-SBD as a next-generation parasite-derived adjuvant for subunit vaccine development.

While this study offers insightful information on the molecular and functional characteristics of *Pf*HSP70-3-SBD as an adjuvant when co-formulated with *Pf*CSP antigen in a murine model, as summarized in the graphical abstract (**Fig. 7**), further evaluation in appropriate *in vivo* challenge models is warranted to establish its protective efficacy and provide a foundation for advancing such adjuvant-like chaperonic domains toward clinical investigation. Furthermore, our findings highlight the adjuvanting potential of *Pf*HSP70-3-SBD, while its intrinsic immunomodulatory and antigenic properties warrant further investigation. Comprehensive characterization of these properties may establish whether *Pf*HSP70-3-SBD can function as both an immunostimulatory and antigenic component, potentially supporting the development of multifunctional vaccine formulations against malaria. Collectively, our findings identify *Pf*HSP70-3-SBD as a parasite-derived component capable of enhancing complementary humoral and cellular immune responses. Beyond its application in malaria vaccination, this antigen-delivery adjuvant approach provides a promising framework for coupling antigen delivery with immune stimulation and may be adaptable to the development of next-generation protein subunit vaccines against a broader range of infectious diseases.

**Fig. 7:**
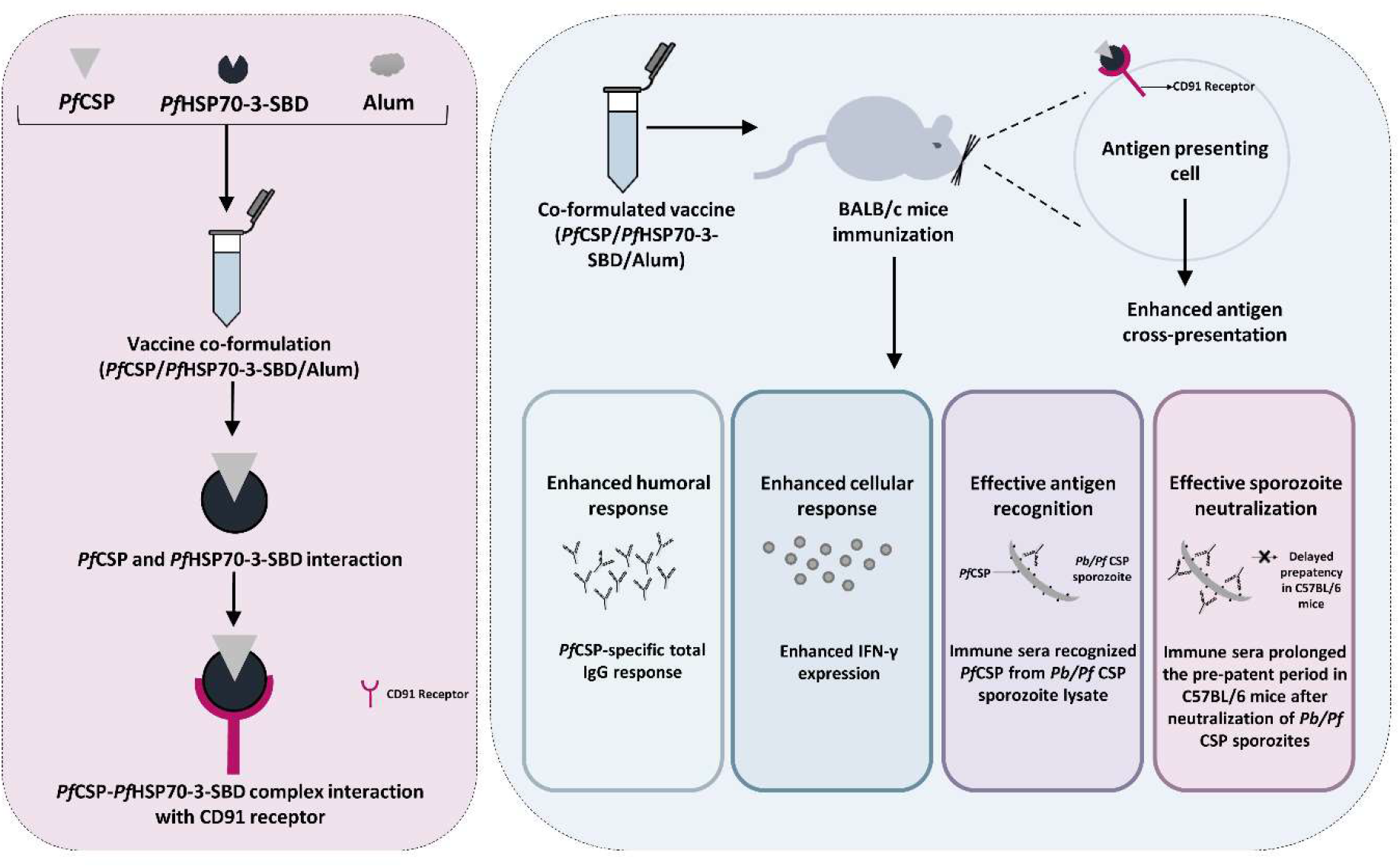
Schematic representation of the evaluation of *Pf*HSP70-3-SBD as an antigen-delivery adjuvant in a *Pf*CSP-based subunit malaria vaccine formulation. *In silico* and *in vitro* characterization demonstrated an interaction between *Pf*HSP70-3-SBD and *Pf*CSP. Furthermore, the predicted *in silico* interaction of the *Pf*CSP-*Pf*HSP70-3-SBD complex with the host CD91 receptor suggests a possible mechanism involving receptor-mediated internalization and subsequent cross-presentation of the associated *Pf*CSP antigen. In BALB/c mice, co-formulation of *Pf*HSP70-3-SBD with *Pf*CSP in an alum-based protein subunit vaccine elicited higher *Pf*CSP-specific total IgG responses and increased IFN-γ expression, indicating the potential of *Pf*HSP70-3-SBD to augment both humoral and cellular immune responses. Furthermore, immunization with *Pf*CSP/*Pf*HSP70-3-SBD/Alum induced antibodies that recognized *Pf*CSP expressed by transgenic *Pb/Pf*CSP sporozoites. In a transgenic sporozoite neutralization assay, this formulation was associated with delayed parasite prepatency in C57BL/6 mice, supporting its functional activity against the pre-erythrocytic stage of the parasite. Collectively, these findings identify *Pf*HSP70-3-SBD as a promising parasite-derived adjuvant candidate for improving the antigen-specific immunogenicity and functional efficacy of *Pf*CSP-based subunit malaria vaccines.

## CONTRIBUTORS

Conceptualization: S.S.

Methodology: S.S., A.R., K.A.K., P.G., C.K.V.

Investigation: P.G., C.K.V., K.G., A.S., S.J., N.J., SA.S.

Visualization: P.G., C.K.V., K.G.

Funding acquisition: S.S.

Supervision: S.S., A.R., K.A.K.

Writing—original draft: S.S., A.R., K.A.K., P.G., C.K.V., K.G.

Writing—review & editing: All authors.

Manuscript was approved by all authors who contributed to it

## DATA SHARING

The primary manuscript and/or the supplementary materials contain information that supports the study’s conclusions and findings. The plasmids generated and/or used in this study are available from S.S. upon reasonable request and the execution of an appropriate Material Transfer Agreement (MTA).

## ACKNOWLEDGEMENTS

We are grateful to Central Laboratory Animal Resources (CLAR), Jawaharlal Nehru University, and the Animal House Facility, University of Hyderabad, for providing facilities to conduct animal experiments. We also thank JNU for providing access to the Turnitin anti-plagiarism software. We are also grateful to the University of Hyderabad for providing an insectary facility. We are also thankful to the Central Instrumentation Facilities (CIF) of Special Centre for Molecular Medicine, JNU. P.G. is supported by DBT-SRF. A.S. is supported by CSIR-UGC JRF. K.G. is a recipient of the PMRF. SA.S. is supported by CSIR-UGC SRF. S.S. is a recipient of the National Bio Scientist Award. This research was supported by funding from the Indian Council of Medical Research (IIRPSG-2024-01-03940) sanctioned to S.S.

