## Supplementary file for "Evolutionarily Conserved *Plasmodium falciparum* HSP70-3 Enhances the Immunogenicity of the Established Malaria Vaccine Antigen Circumsporozoite Protein through Cross-Presentation"

#### SUPPLEMENTARY FIGURES

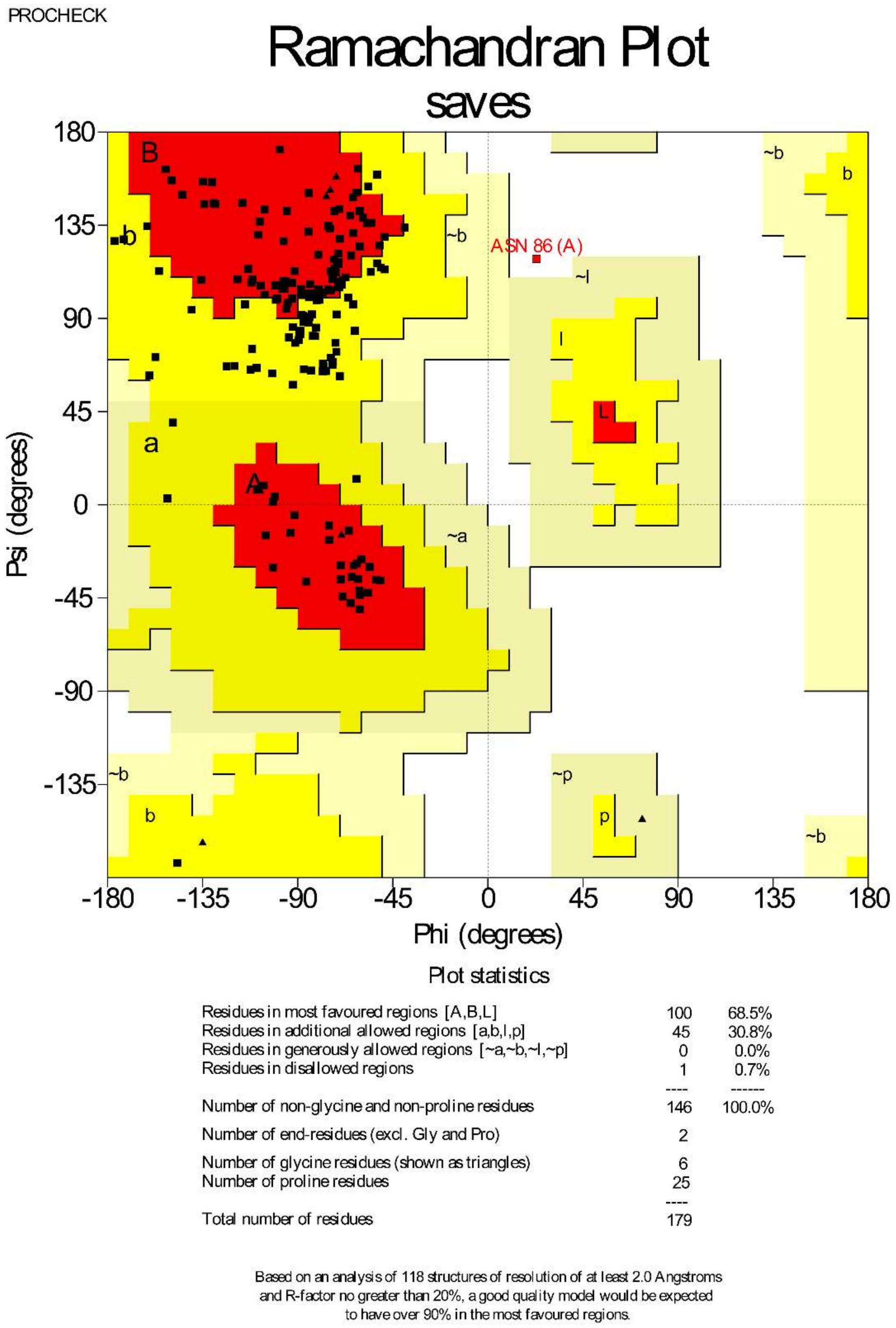


**Fig. S1: PROCHECK Ramachandran plot validation for *Pf*CSP model.** Analysis indicates 68.5% of residues in most favoured, 30.8% in allowed, 0.0% in generously allowed, and 0.7% in disallowed regions, demonstrating acceptable backbone geometry for core structural domains.

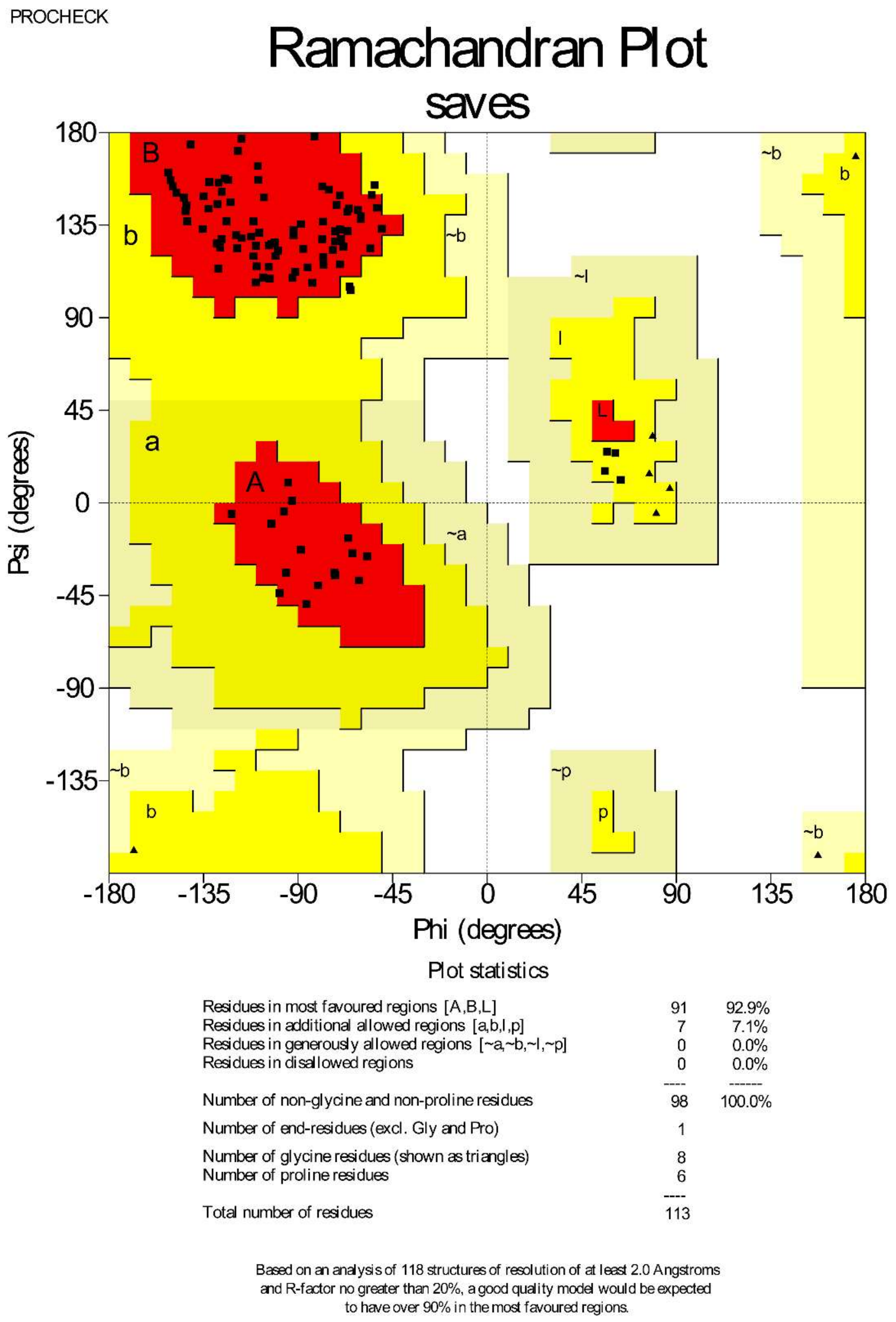


**Fig. S2: PROCHECK Ramachandran plot validation for *Pf*HSP70-3-SBD model.** Analysis indicates 92.9% of residues in most favoured regions, 7.1% in allowed regions, and 0.0% in disallowed regions, confirming high stereochemical quality suitable for docking studies.

### SUPPLEMENTARY TABLES

**Table S1: Amino acid sequence details of the *Pf*HSP70-3-SBD and *Pf*CSP regions used in this study**

| S.No. | Protein | Sequence |
|---|---|---|
| 1. | *Pf*HSP70-3-SBD | DVIPLSLGIETLGGVFTKLINRNTTIPTKKSQIFSTAADNQTQVSIKVFQGEREMASDNKLLGS FDLVGIPPAPRGVPQIEVTFDVDANAIINISAIDKMTNKKQQITIQSSG |
| 2. | *Pf*CSP | DPNANPNANPNANPNANPNANPNANPNANPNANPNANPNANPNANPNANPNANPNANPNANPNA NPNANPNANPNANPNKNNQGNGQGHNMPNDPNRNVDENANANSAVKNNNNEEPSDKHIK EYLNKIQNSLSTEWSPCSVTCGNGIQVRIKPGSANKPKDELDYANDIEKKICKMEKCSSV |

**Table S2: Details of primer sequences used for cytokine analysis by qPCR real-time analysis**

| S.No. | Primer | Sequence |
|---|---|---|
| 1. | GAPDH forward primer | 5’CATCACTGCCACCCAGAAGACTG 3’ |
| 2. | GAPDH reverse primer | 5’ATGCCAGTGAGCTTCCCGTTCAG 3’ |
| 3. | IFN-γ forward primer | 5 ′ AAAGAGATAATCTGGCTCTGC 3′ |
| 4. | IFN-γ reverse primer | 5 ′ GCTCTGAGACAATGAACGCT 3′ |

**Table S3: Analysis of pre-patency in C57BL/6 mice following intravenous challenge with transgenic *Pb/Pf* CSP sporozoites after neutralization**

| S.No. | Group (dilution of immune serum or mAb) | Number of sporozoites per mouse | Mice positive for parasitemia/ mice injected | Pre-patent period (day) | Long-term outcome (2 months post-challenge) |
|---|---|---|---|---|---|
| 1. | Control (1:10 dilution) | 5 x $10^3$ | 5/5 | 3.5 | All mice died |
| 2. | *Pf*CSP/*Pf*HSP70-3-SBD/Alum (1:10 dilution) | 5 x $10^3$ | 2/5 | 5.5 | |
| | | | 1/5 | 6.5 | |
| | | | 2/5 | No patency till day 12 | 2/5 mice survived without detectable parasitemia |
| 3. | mAb 2A10 (1:50 dilution) | 5 x $10^3$ | 0/5 | No patency till day 12 | 5/5 mice survived without detectable parasitemia |

Pre-patency is defined as the time required for appearance of blood-stage infection following infection of mice with infectious sporozoites.
